# Gut Bacteria Prime Host Antibody Responses Against Ingested Dietary Fiber Glycans

**DOI:** 10.64898/2026.08.07.743361

**Authors:** Giovanni Vega, Carolina Agudelo, Meghan E. Graham, Elyza A. Do, Jasmin Akter, Rashidul Haque, Nicolette Hernandez-Kaempf, Khurshid R. Iranpur, Abigail Renfro, Ansel Hsiao, Ashley R. Wolf, Michael L. Patnode

## Abstract

Protein and glycan antigens synthesized by gut microbes stimulate circulating and secreted antibody production. Despite continuous exposure of hosts to the plant glycans that constitute dietary fiber, it remains unclear whether these foreign structures induce mucosal immune responses. We report that humans and mice generate antibodies specific for common glycans in plant foods. Oral exposure to individual fiber types induced T cell-independent, glycan-specific IgM and IgA. The induction of anti-fiber antibodies required colonization by particular microbes, since germ-free mice and mice harboring representatives of several bacterial phyla failed to respond. The OMM12 model community was sufficient to rescue antibody induction, and dietary fiber glycans were detected on the surfaces of OMM12 microbes, suggesting a route by which bacteria trigger anti-fiber immune responses. Our results reveal a direct impact of dietary fiber on the adaptive immune system with implications for host control of fiber breakdown by bacteria in the gut lumen.

## Main

Dietary fiber is an abundant component of nearly all mammalian diets and is composed of ingested carbohydrates that are not susceptible to degradation by host enzymes. In typical human diets, fiber is composed primarily of high molecular weight plant cell wall polysaccharides, which serve as the major energy and carbon source for the most abundant gut bacterial taxa^1^. Mechanisms by which the host can modulate microbial metabolism may provide evolutionary advantages^2,3^, yet it is unclear whether, or how, microbial access to carbohydrates in the gut lumen is subject to host control. Although plant cell wall polysaccharides have not previously been considered immunogenic in their ingested forms, antibodies targeting these glycans could have an outsized impact on microbial food webs in the gut. Immune responses against microbial polysaccharides are crucial for host protection and form the basis of several existing vaccines^4,5^. Antibodies against capsular polysaccharides and lipopolysaccharides are typically produced through a T cell-independent pathway that results in strong IgM responses^6,7^. Additionally, association of these glycans with intact bacteria or a carrier protein can lead to an enhanced response, including long-term immunity and antibody class-switching^8,9^.

Gut bacteria bind with high affinity and specificity to dietary fiber glycans^10^, raising the possibility that microbial cell surfaces might be coated with fiber glycans *in vivo* and that these bacteria initiate bystander immune responses against plant polysaccharides. We reasoned that fiber glycans have the potential to trigger antibody responses in mammals because they are (i) physically associated with bacteria that can induce co-stimulation of B cells via innate immune receptors, (ii) abundant in the intestinal lumen offering the chance for repeated exposure throughout life, and (iii) distinct from host glycans suggesting that circulating naive B cell clones might recognize these non-self structures. Here, we employed screening approaches using glycan coated beads^11^ and detected antibodies specific for dietary fiber produced during steady state in humans and mice. We found that dietary fiber supplementation regimens modulate anti-fiber antibodies on short timescales. Gnotobiotic mice colonized with individual strains, model consortia, and microbial communities of native complexity revealed that exposure to specific microbes is required to sensitize mice to dietary fiber. Together, these results uncover a new class of immune responses directed against diet and raise the prospect of an inducible mode of host control over fiber-degrading bacteria in the gut.

## Results

### Humans produce circulating IgM and secreted IgA specific for dietary fiber glycans

To determine whether humans produce antibodies that recognize dietary fiber glycans, we measured IgM reactivity in serum samples from adults (n=17, 20-41 years old) and their children (n=19, 3-4 years old) at two timepoints one week apart **(Fig. 1a)**. We used multi-color flow cytometry to analyze a pooled library of microscopic magnetic beads coated with distinct glycan preparations representing the major classes of plant cell wall polysaccharides and tagged with unique fluorescent barcodes^10^. Each glycan preparation has been characterized by mass spectrometry with respect to monosaccharide composition and glycosidic linkages present^10^. This screen revealed significant IgM binding to 30 bead types, including those coated with plant polysaccharides commonly found in human diets such as cell wall hemicelluloses (xylans and galactomannans) and multiple components of pectins **(Fig. 1b and Table S1a)**. Serum collected from the same donors one week apart showed that many anti-fiber IgM profiles were consistent **(Table S1a)**, including 15 fiber types displaying strong correlations between the two timepoints across donors (Spearman’s rho > 0.7, **Fig. 1b and Table S1a**). IgM reactivity was higher in children compared to adults for 6 out of the 56 bead types in the library, despite the generally observed increase in IgM levels with increasing age^12,13^. We did not observe significant correlations between the antibody profiles of mothers and their own children, with samples instead clustering by age group. Based on the known correspondence between IgA specificity in breast milk and at other mucosal sites including the intestine^14,15^, we tested pooled human colostrum and observed IgA binding to arabinogalactan A, galactan, oat-beta glucan, psyllium, and rhamnogalacturonan K **(Fig. 1c and Table S1b)**. These data demonstrate that humans produce circulating fiber-specific IgM under steady state conditions and also secrete anti-fiber IgA (sIgA) with similar glycan specificities.

**Fig. 1.**
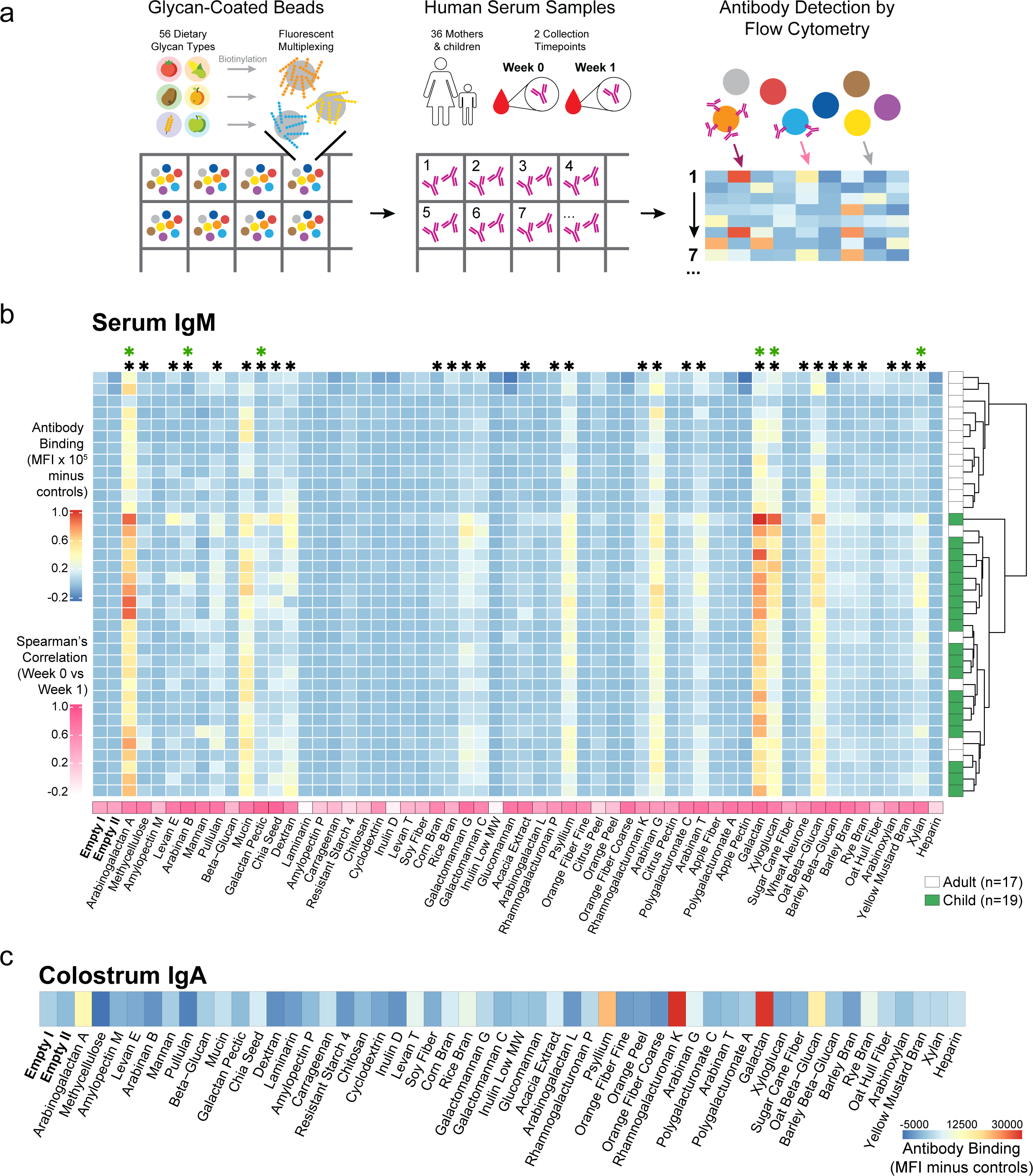
Adults and children produce circulating anti-fiber IgM against dietary glycans. (a) Schematic illustrating serum collected from mother-child pairs at two timepoints (one week apart). Serum was screened by flow cytometry for circulating anti-fiber IgM using a library of magnetic beads coated with 56 distinct glycan preparations. (b) Heatmap showing human serum IgM binding to glycan-coated beads (blue to red color scale). The plotted values show the average signal of week 0 and week 1 collections (black *, Paired t test, q < 0.1, FC >= 1.5 at both timepoints versus empty beads; green *, Student’s t test, q < 0.05, FC >= 1.25 at both timepoints between adults and children). The rows represent individual donors and samples are clustered (Ward’s minimum variance) by binding profile (rightmost column indicates sample age group; adults, white; children, green). The bottom row (white to pink color scale) shows Spearman’s correlations between the two timepoints. (c) Human colostrum was tested for the presence of anti-fiber IgA using the glycan-coated bead library. The row represents the average of 3 technical replicates. Heatmap values are corrected by subtracting the average of 3 secondary-only controls and the average of 2 empty bead populations.

### Ingestion of fiber induces glycan-specific anti-fiber IgM and IgA

To determine whether oral exposure to dietary fiber influences the levels of fiber-specific antibodies, we compared circulating anti-fiber antibodies in C57BL/6 mice bred for four generations on either a standard mouse chow control diet or a diet with low levels of microbially accessible fibers (“low fiber diet”). The low fiber diet led to significantly lower circulating IgM reactivity against plant glycans including arabinans and xylans **(Fig. 2a and Table S2a)**, which are ubiquitous cell wall polysaccharides that are present in the corn and wheat ingredients in standard mouse chow. Mouse IgM reactivity was similar to that observed in humans **(Fig. 1b)**, consistent with the abundance of these structures in both mouse and human diets. Serum IgG and IgA bound a small and distinct set of bead types, and only the binding to levan and porcine mucin were significantly reduced in the low fiber diet group **(Extended Data Fig. 1a-b and Table S3a-b)**. Together, these data indicate that consumption of a diet containing abundant fiber induces antibody responses against a broad range of plant polysaccharides.

**Fig. 2.**
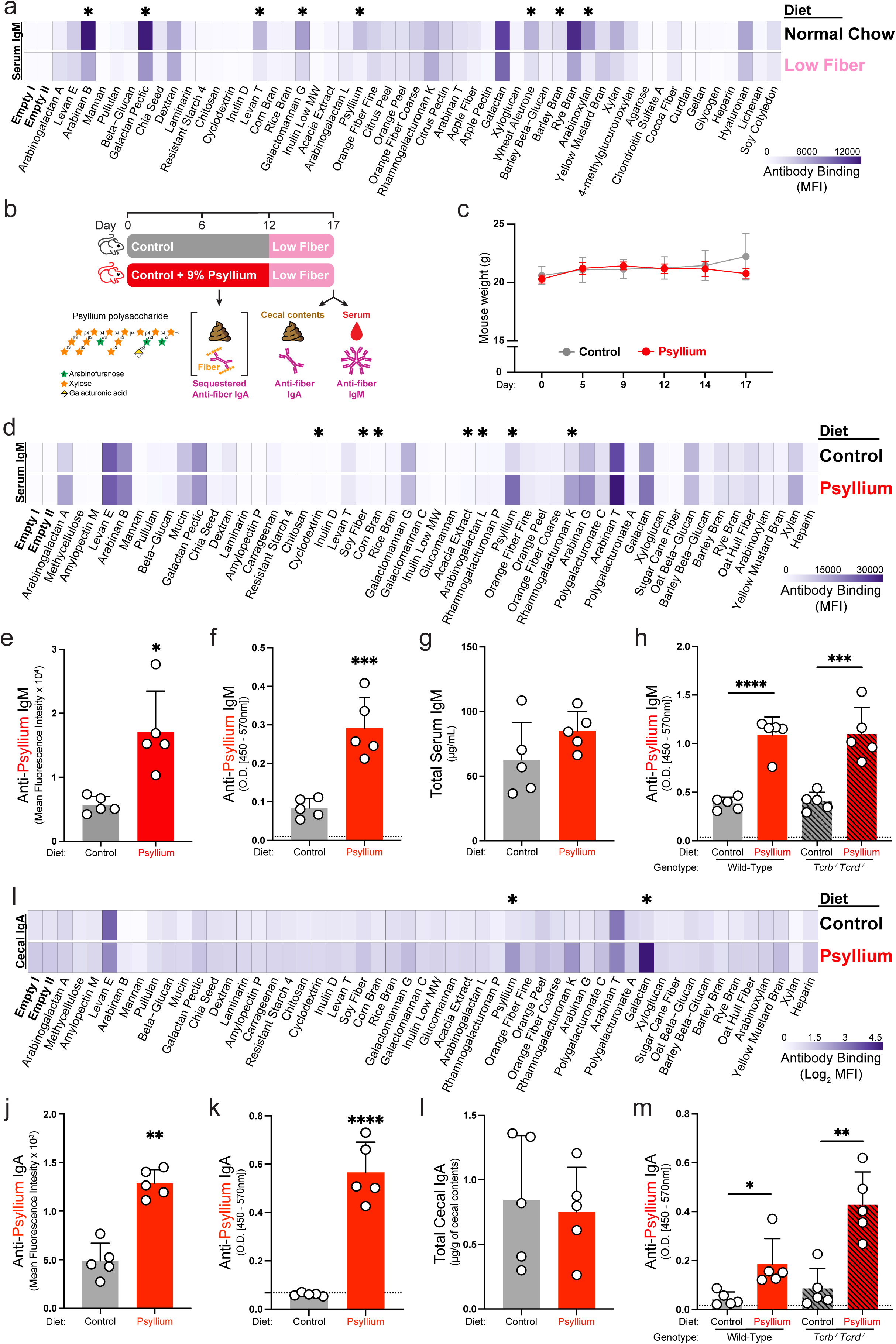
Fiber supplementation induces circulating and secreted fiber-specific antibodies. (a) Anti-fiber IgM was measured in serum from C57BL/6 mice eating normal chow (2920X, Envigo) or a low fiber diet (TD.130343, Envigo) by flow cytometry using a glycan-coated bead library. Each row shows the average of 7-8 mice. Results are representative of 2 independent experiments. (b) Schematic illustrating C57BL/6 mice fed a control or 9.2% w/w psyllium-supplemented diet for 12 days followed by a low fiber diet for 5 days. Brackets depict dietary fiber in the cecal contents sequestering anti-fiber antibodies and blocking their detection. Cecal contents and serum were collected on day 17. (c) Mouse weights throughout a representative feeding experiment. (d) Anti-fiber IgM in serum on day 17. Each row represents the average of 5 mice. (e) Individual values for anti-psyllium IgM in serum on day 17. (f) ELISA for anti-psyllium IgM in serum on day 17 (Student’s t test, ***p < 0.001). (g) Quantification of total serum IgM on day 17. Results for (c - g) are representative of >3 independent experiments. (h) Anti-psyllium IgM in serum from wild-type or *Tcrb^-/-^Tcrd^-/-^* C57BL/6 mice on day 17 of the experimental feeding protocol shown in b (Student’s t test, ***p < 0.001 and ****p < 0.0001). Results are representative of 3 independent experiments. (i) Anti-fiber IgA in cecal contents on day 17. Each row represents the average of 5 mice. (j) Individual values for anti-psyllium IgA in cecal contents on day 17. (k) ELISA for anti-psyllium IgA in cecal contents on day 17 (Student’s t test, ****p < 0.0001). (l) Quantification of total cecal IgA on day 17. Results for (i-l) are representative of 3 independent experiments. (m) Anti-psyllium IgA in cecal contents from wild-type or *Tcrb^-/-^Tcrd^-/-^*C57BL/6 mice on day 17 of the experimental feeding protocol shown in B (Student’s t test, *p < 0.05 and **p < 0.01). Results are representative of 3 independent experiments. For heatmaps, the plotted values are normalized by subtracting the average of 3 secondary-only controls. When linear values are shown (a and d), the average of 2 empty bead populations are subtracted (*, Student’s t test, q < 0.05, FC >= 1.5). (i) The plotted values are normalized by subtracting the average of 3 secondary-only controls and log2 transformed (*, Student’s t test and q < 0.05). (e and j) * q < 0.05 and ** q < 0.01. Dotted lines, background signal (average of 3 secondary-only controls); bars, mean + SD.

The observed dependence of anti-fiber antibodies on diet suggested that exposure to a variety of diverse plant polysaccharides was responsible for distinct antibody responses. However, this could alternatively indicate the presence of polyreactive antibodies that broadly recognize many unrelated glycan structures. To determine whether antibody responses to fiber are glycan-specific, we fed C57BL/6 mice food-grade psyllium husk, which contains a beta-1,4-xylose backbone densely substituted with xylose and arabinose side chains variably capped with terminal galacturonic acid. Psyllium was selected because (i) it is one of the fiber supplements most commonly consumed by humans^16–18^, (ii) anti-psyllium IgM exhibited a large fold-change in mice fed a control versus low fiber diet **(Fig. 2a)**, and (iii) humans produce anti-psyllium IgM and IgA **(Fig. 1b-c)**. Mice were maintained for 12 days on either a control mouse chow diet or that same diet supplemented with 9.2% w/w psyllium, followed by administration of a low fiber diet for 5 days to limit sequestration of secreted anti-psyllium IgA by structurally similar xylans in normal chow **(Fig. 2b)**. No difference in mouse weight was observed between the two groups during feeding **(Fig. 2c)**. Circulating anti-psyllium IgM in mice fed a psyllium diet was 3-fold higher than in the control group **(Fig. 2d-e and Table S2b)**. Higher levels of antibodies targeting rhamnogalacturonan from karaya (2.4-fold) were detected in these mice, potentially reflecting the glycan epitopes that this fiber shares with psyllium^10^. The antibodies induced by psyllium supplementation were notably glycan-specific, as there was no detectable increase for fiber types with only subtle differences from psyllium in glycan structure (more sparsely substituted wheat arabinoxylan and beechwood xylan) and the highest signal for other unrelated fiber types was 7.1-fold lower than for psyllium. Measurement of anti-psyllium IgM by ELISA produced similar results, with mice eating a psyllium diet exhibiting 3.5-fold higher levels than control animals **(Fig. 2f)**. Total serum IgM in the two diet groups was comparable **(Fig. 2g)**, indicating that the anti-psyllium IgM induction reflects a change in antibody repertoire rather than a broad increase in IgM abundance.

Antibodies that recognize glycans and other non-protein antigens are commonly produced via T cell-independent pathways^6,7^. To determine whether the induction of fiber-specific antibodies is dependent on T cells, we fed C57BL/6 wild-type and *Tcrb^-/-^Tcrd^-/-^* mice psyllium for 12 days followed by a low fiber diet for 5 days, as above. Two days prior to fiber feeding, all groups of mice exhibited comparable circulating anti-fiber IgM profiles, with the exception of reactivity against coarse orange fiber, arabinan G, and arabinan T, which were elevated in T cell-deficient mice **(Extended Data Fig. 2a-b and Table S4a)**. Upon psyllium feeding, both wild-type and T cell-deficient mice exhibited a >2-fold increase in circulating anti-psyllium IgM **(Fig. 2h, Extended Data Fig. 2c-d, and Table S4b)**, indicating that T cell help is not required for the induction of circulating anti-psyllium IgM.

The oral route of host exposure to dietary fiber led us to ask whether a secreted IgA response accompanies the circulating IgM response against ingested plant glycans. Mice fed psyllium produced 2.6-fold higher levels of anti-psyllium IgA in cecal contents compared to mice eating the control diet, as measured by flow cytometry **(Fig. 2i-j and Table S2c)**. These mice also exhibited higher levels of anti-galactan IgA, but the limited galactose detected in psyllium glycans^10^ suggests that this represents a response to a microbial or alternate dietary antigen. ELISA confirmed the induction of anti-psyllium IgA in mice fed psyllium (9.3-fold induction; **Fig. 2k**). Psyllium did not increase total secreted IgA levels **(Fig. 2l)** and induction of anti-psyllium IgA was T cell-independent (4.2-fold increase in wild-type mice and 5-fold increase in *Tcrb^-/-^ Tcrd^-/-^*mice) **(Fig. 2m)**. Taken together, these results demonstrate that ingestion of dietary fiber stimulates the production of circulating anti-fiber IgM and secreted anti-fiber IgA independent of T cell help.

### Induction of anti-fiber antibodies requires exposure to gut microbes

Ingested plant cell wall polysaccharides are not regarded as immunogenic, and there are no known innate recognition pathways involved in the detection of these glycans by B cells. However, we reasoned that gut bacterial products could provide co-stimulation during B cell responses to dietary plant glycans, especially if those bacteria physically associate with fiber^10^. The impact of the gut microbiota on the baseline levels of anti-fiber IgM was modest, with reactivity against 10 fiber types reduced in germ-free mice, including fructans (levan E, levan T, inulin low MW), beta-glucans (low viscosity beta-glucan, oat beta-glucan), and arabinoxylans (low viscosity arabinoxylan, corn bran, rye bran, oat hull fiber) **(Extended Data Fig. 3a and Table S5a)**. To determine whether microbes are required for the induction of anti-fiber antibodies, germ-free and conventionally raised mice were fed a control or psyllium-supplemented diet for 12 days. Germ-free mice failed to produce circulating anti-psyllium IgM in response to psyllium feeding **(Fig. 3a)**. Baseline measurements of anti-fiber IgM analyzed 4 days before the fiber feedings confirmed that induction was restricted to conventionally raised mice fed psyllium and that psyllium did not influence the abundance of antibodies against unrelated glycans, such as galactomannan **(Fig. 3b-c, Extended Data Fig. 3b and Table S5b)**. In conventionally raised Swiss Webster (SW) mice, increased circulating anti-psyllium IgM was observed in response to psyllium feeding **(Extended Data Fig. 4a-f and Table S6a-b)**, and the high variance and lack of a detectable IgA response is consistent with the more variable antibody responses in this genetically diverse outbred strain^19,20^. Like C57BL/6 mice, germ-free SW mice failed to generate anti-psyllium antibodies after psyllium feedings **(Extended Data Fig. 4g-i and Table S6c)**. We also confirmed that the high-temperature autoclave protocol used for sterilizing germ-free food and bedding did not abrogate the anti-psyllium response in conventionally raised C57BL/6 mice **(Extended Data Fig. 5a-c and Table S7)**. Thus, microbial exposure is required for the diet-induced production of circulating anti-fiber antibodies in multiple genetic backgrounds.

**Fig. 3.**
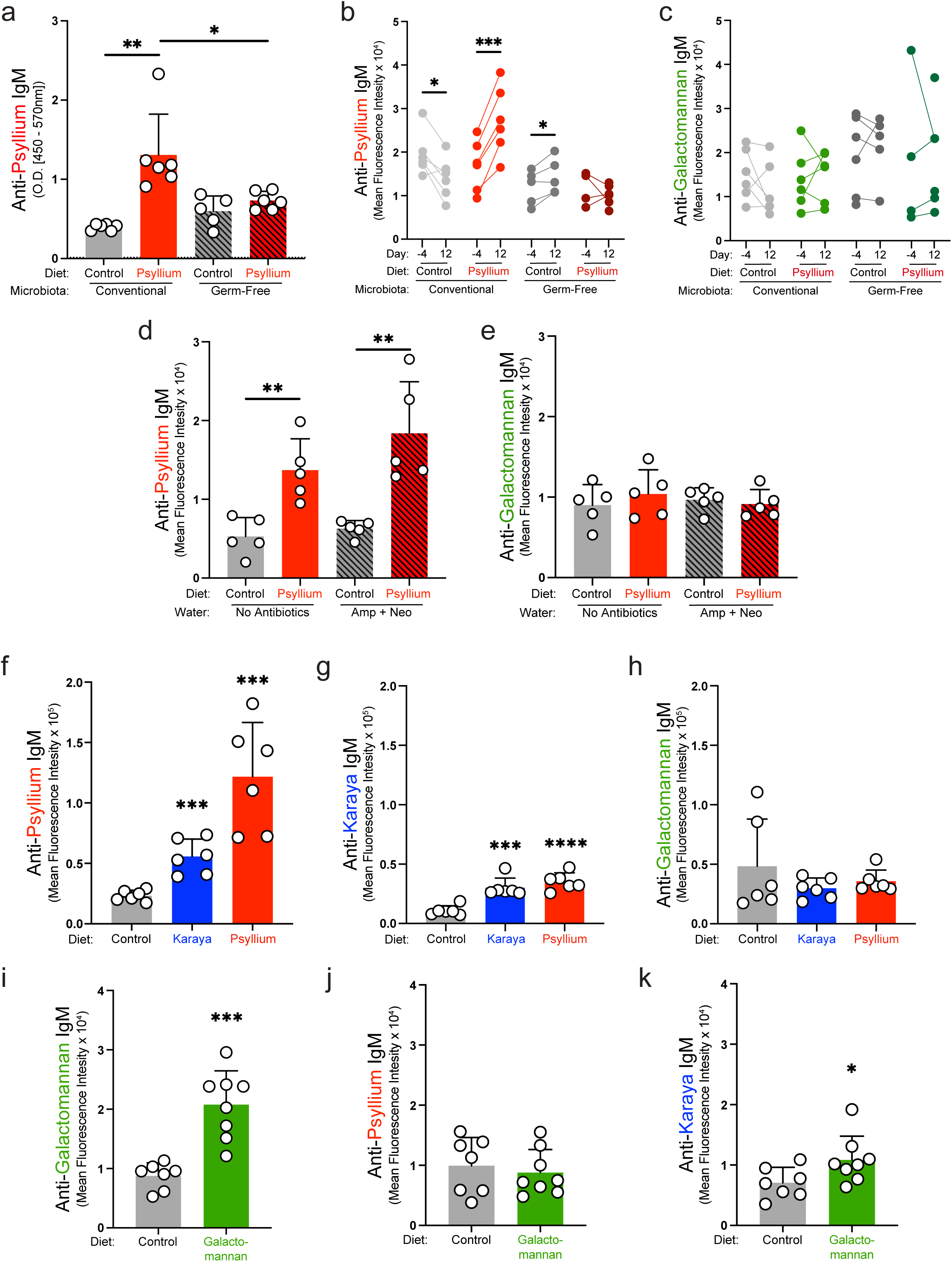
Microbes are required for the induction of anti-fiber antibodies. (a) Anti-psyllium IgM in serum from conventionally raised or germ-free C57BL/6 mice eating a control or 9.2% w/w psyllium-supplemented diet for 12 days (Student’s t test, *p < 0.05 and **p < 0.01). Results are representative of >3 independent experiments. (b and c) Serum from conventionally raised or germ-free C57BL/6 mice collected before (day -4) and after eating a control or psyllium-supplemented diet (day 12) was tested using a glycan-coated bead library to measure baseline differences of circulating anti-psyllium IgM (b) and anti-galactomannan IgM (c). Dots represent individual mice, with lines connecting paired samples, and unpaired samples were plotted with no connecting line (Paired t test, *p < 0.05 and ***p < 0.001). (d) Anti-psyllium IgM and (e) anti-galactomannan IgM in serum from conventionally raised C57BL/6 mice, with or without continuous antibiotic treatment, eating a control or 9.2% w/w psyllium-supplemented diet for 12 days (Student’s t test, **p < 0.01). (f - h) Anti-psylium IgM (f), anti-karaya IgM (g), or anti-galactomannan IgM (h) in serum from C57BL/6 mice eating a control, 9.2% w/w psyllium-, or 9.2% w/w karaya-supplemented diet for 12 days (Student’s t test, ***p < 0.001 and ****p < 0.0001). (I - K) Anti-galactomannan IgM (i), anti-psyllium IgM (j), or anti-karaya IgM (k) in serum from C57BL/6 mice eating a control or 9.2% w/w galactomannan-supplemented diet for 11 days (Student’s t test, *p < 0.05 and ***p < 0.001). Results are representative of 3 independent experiments. Plotted MFI values are normalized by subtracting the average of 3 secondary-only controls and the average of 2 empty bead populations. Dotted lines, background signal (average of 3 secondary-only controls); bars, mean + SD.

To determine the impact of fiber-induced changes to the gut microbiota on the induction of fiber-specific antibodies, antibiotics (ampicillin and neomycin) were added to the drinking water of conventionally raised mice (starting 4 days before the psyllium feeding). The treatment depleted bacterial biomass as assessed by measuring fecal DNA on day -4 through 12 **(Extended Data Fig. 6a)**. Mice exhibited an induction of anti-psyllium antibodies regardless of antibiotic treatment (2.6-fold increase versus 3-fold increase; **Fig. 3d, Extended Data Fig. 6b-c, and Table S8**). The induction of anti-fiber antibodies was specific to psyllium as neither psyllium feeding nor antibiotic treatment influenced anti-galactomannan IgM levels **(Fig. 3e)**. These results are consistent with a limited role for microbes during fiber supplementation and suggest microbial exposure before dietary intervention is necessary for generating anti-fiber responses.

Although microbial changes induced by psyllium did not appear to be involved in the induction of circulating anti-psyllium IgM, fiber-induced changes in microbiota composition could be involved in IgM induction against other fiber types that are more readily degraded by the gut microbiota, such as karaya gum, rich in rhamnogalacturonan II^21^. Mice were fed a control, psyllium-supplemented, or karaya gum-supplemented diet for 12 days and the fecal microbiota composition was analyzed by performing 16S rDNA sequencing^22^. As expected, both diets produced alterations in microbiota composition that were dependent on the fiber type consumed, with psyllium increasing the relative abundance of *Akkermansia*, *Parasutterella*, and *Bacteroides* within the first 5 days **(Extended Data Fig. 7a-b and Table S9)**, consistent with previous studies^23,24^. Since mice in our facility typically lack representatives of the *Proteobacteria* phylum, all mice were additionally inoculated with commensal *Escherichia coli* on day 5. By day 12, psyllium continued to induce distinct microbiota compositions **(Extended Data Fig. 7c)**. Both diets resulted in the production of anti-fiber antibodies compared to the control diet, with psyllium inducing a 5.2-fold increase in anti-psyllium IgM and karaya inducing a 2.8-fold increase in anti-karaya IgM **(Fig. 3f-g)**. The induction of anti-psyllium IgM by karaya (2.4-fold increase) and anti-karaya IgM by psyllium (3.3-fold increase) is likely due to presence of similar glycan antigens in both fiber types^10^. Consistent with our previous feeding results, neither diet increased anti-fiber antibodies against galactomannan **(Fig. 3h)**.

To test an additional readily fermentable fiber with a distinct monosaccharide composition, we fed mice a diet containing 9.2% w/w guar gum (galactomannan G) composed of a beta-1,4-mannan backbone with alpha-1,6-galactose side chains^25,26^. We observed a 2.4-fold increase in circulating anti-galactomannan IgM in mice eating a galactomannan-containing diet compared to the control diet **(Fig. 3i, Extended Data Fig. 8a, and Table S10)**. The induction of anti-galactomannan was validated by ELISA **(Extended Data Fig. 8b)** and was not due to a global increase in total serum IgM **(Extended Data Fig. 8c)**. As expected based on the lack of induction of anti-galactomannan antibodies when mice were fed psyllium **(Fig. 2d)**, galactomannan did not induce anti-psylium antibodies **(Fig. 3j)**. We detected modestly higher anti-karaya antibodies (1.5-fold) in mice fed galactomannan, which may be due to the abundant galactose side chains in karaya **(Fig. 3k)**. These data, along with the results of antibiotic treatment, argue against the notion that a single stereotyped microbiota alteration during fiber exposure is responsible for the induction of fiber-specific antibodies.

To assess which microbial taxa were sufficient for enabling anti-fiber immune responses, germ-free mice were colonized with minimal communities containing *Bacteroides ovatus* (ATCC-8483) in combination with (i) *E. coli* (TSDC17.2), (ii) *E. coli* (TSDC17.2) plus *Akkermansia muciniphila* (NSD001), or (iii) *Bacteroides ovatus* (WH514), and fed a control or psyllium-containing diet. None of these highly simplified yet phylogenetically diverse collections of taxa were sufficient to permit significant antibody responses against ingested psyllium **(Extended Data Fig. 9a-d and Table S11a-b)**. Together, these results indicate that the induction of circulating anti-fiber antibodies results from immune activation by ingested dietary fiber glycans in a manner dependent on particular microbial taxa prior to oral fiber exposure.

### Culturable commensal gut bacteria enable the induction of fiber-specific antibodies

While minimal bacterial communities appeared to lack the necessary signals to sensitize mice to produce anti-fiber antibodies, we wondered whether a broader collection of culturable members of the endogenous mouse microbiota was sufficient. Fecal bacteria from mice eating a low fiber diet were plated on BHI blood agar and then combined to create a pooled “culture collection”. Male and female germ-free mice colonized with this culture collection were bred and their progeny were fed a control or psyllium-supplemented diet for 12 days followed by a low fiber diet for 5 days. Psyllium feeding induced 1.4-fold higher anti-psyllium IgM levels in the serum of culture collection colonized mice **(Fig. 4a, Extended Data Fig. 10a-b, and Table S12a)**. In a parallel arm of the experiment, fecal pellets collected from these mice 2 weeks prior to psyllium feeding were used to colonize age-matched germ-free mice. The mice colonized as adults also exhibited an increase in anti-psyllium IgM (1.9-fold) compared to mice eating a control diet **(Fig. 4a, Extended Data Fig. 10a-b, and Table S12a)**. Neither group of mice colonized with the culture collection fully recapitulated the anti-psyllium IgA response seen in conventionally raised animals **(Fig. 4b)**. These results reveal that a complex community of culturable members can sensitize mice to generate an IgM response against ingested plant glycans, whether colonized at birth or as adults.

**Fig. 4.**
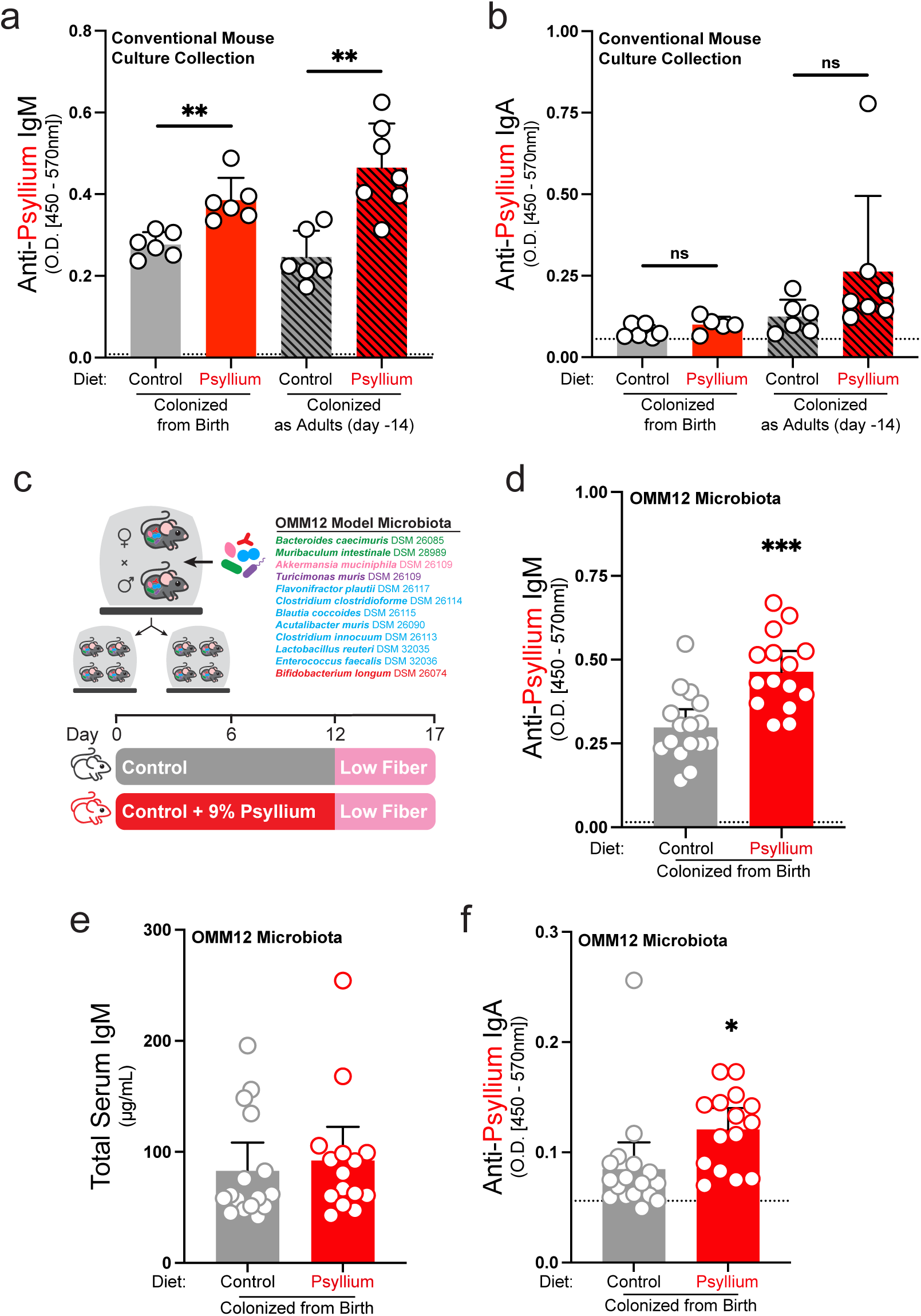
Mouse-derived communities of culturable isolates sensitize mice to psyllium. (a and b) Anti-psyllium IgM in serum (a) and anti-psyllium IgA in cecal contents (b) from germ-free C57BL/6 mice colonized with a conventionally raised mouse culture collection from birth or as adults and fed control or 9.2% w/w psyllium-supplemented diet for 12 days followed by a low fiber diet for 5 days (Student’s t test, **p < 0.01). (c) Schematic showing the OMM12 model community experimental design. (d) Anti-psyllium IgM in serum (Student’s t test, ***p < 0.001). Results are representative of 2 independent experiments. (e) Total IgM in serum. (f) Anti-psyllium IgA in cecal contents (Student’s t test, *p < 0.05). Dotted lines, background signal (average of 3 secondary-only controls); bars, mean + SD or mean + 95% confidence interval (d - f).

A fully defined gut microbial community offers opportunities to understand the mechanisms driving antibody induction. The OMM12 (Oligo-Mouse-Microbiota) model community is reproducible for multiple generations, encompasses broad phylogenetic diversity, and has been fully sequenced^27^ **(Fig. 4c)**. We colonized C57BL/6 mice with the OMM12 community and fed their progeny a control or psyllium-containing diet for 12 days followed by a low fiber diet for 5 days. The mice fed psyllium exhibited a 1.6-fold increase in circulating anti-psyllium IgM **(Fig. 4d, Extended Data Fig. 10c-d, and Table S12b)**, and this was not due to a global increase in circulating IgM **(Fig. 4e)**. In contrast to the mice colonized with the complete culture collection, we detected a 1.4-fold increase in anti-psyllium IgA as a result of psyllium feeding **(Fig. 4f)**, despite the reduced IgA^+^ plasma cell numbers in mice harboring OMM12 strains^28^. Together, these results show that a fully defined, simplified microbial community sensitizes germ-free mice to produce anti-fiber antibodies in response to a fiber-supplemented diet.

### Monoclonal antibodies specific for diverse plant glycans bind the surfaces of gut microbes

Human gut strains from the *Bacteroidales* order adhere to particles covered with dietary fiber glycans with high affinity both *in vitro* and in the gut lumen^10^, raising the possibility that bacteria initiate bystander immune responses against fiber glycans on their surfaces. To comprehensively map the fiber glycans bound to microbial cell surfaces in the intestines of mice, we used a collection of 144 monoclonal antibodies specific for plant polysaccharides. These antibodies were generated by immunizing mice with glycans, carrier proteins, and microbial adjuvants via repeated intraperitoneal injection^29^ and recognize a wide breadth of epitopes found in hemicellulose and pectin (including galactans, arabinans, mannans, xylans, galacturonans, and glucans). To first assess the specificities of these antibodies for our library of fiber glycans with defined monosaccharide and linkage composition, we incubated each antibody with the bead library and quantified binding by flow cytometry **(Fig. 5a)**. Hierarchical clustering revealed that beads coated with fibers from similar sources such as oat/wheat/barely (group I), apples (group II), and citrus (groups III and IV) exhibited similar antibody binding profiles **(Fig. 5b and Table S13)**. The discrete specificities represented in this collection of monoclonal antibodies also allowed us to detect subtle distinctions in multiple glycans from different sources, including arabinans (from sugar beet, ghatti gum, and tagaranth), arabinogalactans (from acacia and larch), galactomannans (from carob and guar), and rhamnogalacturonans (from potato and karaya). Thus, this antibody collection detects a wide range of dietary plant polysaccharides and can distinguish between epitopes with subtle structural differences.

**Fig. 5.**
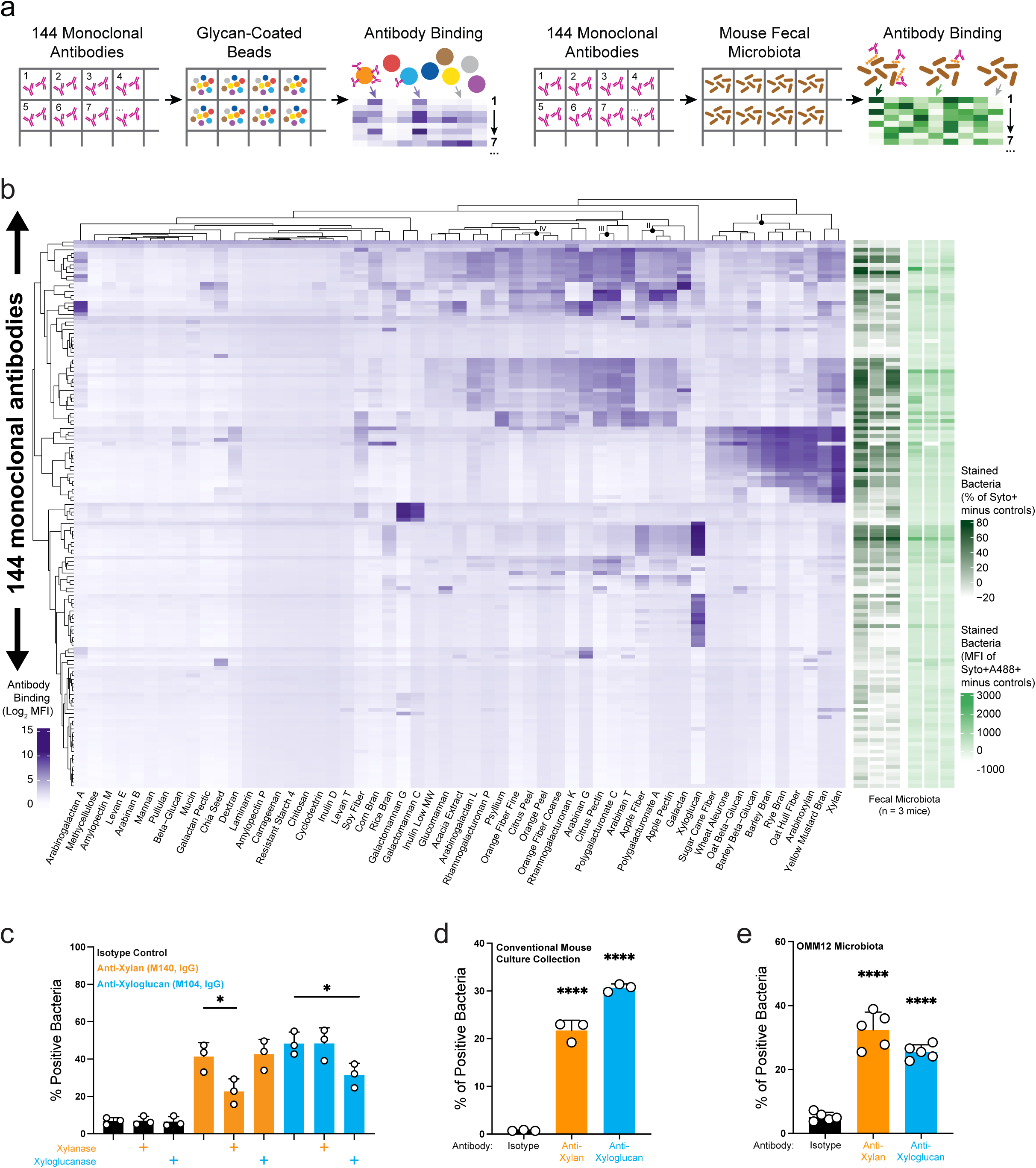
Plant glycans are detected on bacterial surfaces. (a) Schematic depicting high-throughput screening of the plant glycan monoclonal antibody library to define carbohydrate epitopes on the fiber bead library and on gut bacterial surfaces. (b) The left heatmap (purple) shows the specificities of monoclonal antibodies using a glycan-coated bead library. The antibodies and bead types are clustered by binding profiles (Ward’s minimum variance). Each row represents the average of 2 technical replicates. The right heatmaps (green) depict antibody binding percentage (dark green, Syto+ only) and mean fluorescence intensity (light green, Syto+A488+ only) of gut microbes isolated from fecal samples (n = 3 mice) and stained with DNA-binding dye. The plotted values are background corrected by subtracting the average of 3 isotype control samples. (c) Enzymes specific for linkages in plant glycans were used to validate anti-xylan (xylanase) and anti-xyloglucan (xyloglucanase) antibody binding from the monoclonal antibody screen. Fecal pellets were collected from mice eating a low fiber diet and treated with either enzyme before antibody staining (Student’s t test, *p < 0.05). (d and e) Anti-xylan (IgG, M140) or anti-xyloglucan (IgG, M104) binding to bacteria in fecal pellets from gnotobiotic mice colonized with the complete culture collection (d) or the defined OMM12 community (e). (Student’s t test, ****p < 0.0001). Bars, mean + SD.

The antibody panel was next used to identify which dietary polysaccharides are bound to the surfaces of microbes collected directly from the gut lumen. Fecal pellets were harvested from mice (n = 3 mice) eating a low fiber diet to decrease the likelihood of free dietary glycans sequestering the monoclonal antibodies and reducing binding to dietary glycans on bacterial surfaces. Of the 144 tested antibodies, 31 bound both a significantly larger fraction than isotype control and also greater than 25% of the microbial community, and their specificities spanned diverse antigens including, arabinan, galactan, rhamnogalacturonan, xylan, and xyloglucan **(Fig. 5b and Table S13)**. To account for antibody staining of microbes that are present at low abundance (less than 25% of the community), we assessed the MFI (geometric mean fluorescence intensity) of this fraction for those that were significantly higher than the respective isotype control. This revealed 17 additional antibodies that strongly bound to microbial populations with low abundances. Overall, the percentage and MFI profiles were largely consistent with the exception of a low-abundance microbial population stained by antibodies recognizing chia seed. Sets of antibodies that shared similar specificity in the bead-binding assay showed discrepant binding to the same gut microbiota sample, which could be due to the sensitivity of some fiber glycan epitopes to microbial processing. Despite the high specificity of many of these antibodies for linkages unique to plants, it is possible that they exhibit cross-reactivity to microbially synthesized glycans with distinct linkage composition. We used glycoside hydrolases specific for beta-1,4-xylan (xylanase) and beta-1,4-xyloglucan (xyloglucanase) to cleave glycans on microbial surfaces before staining. Treatment with each enzyme reduced the binding of the corresponding antibody **(Fig. 5c)**, indicating the presence of stereotypical plant glycan structures. Xylan (21% binding) and xyloglucan (30% binding) were also detected on microbes in fecal pellets from mice colonized with the complete mouse microbiota culture collection **(Fig. 5d)**. In addition, significant populations of OMM12 gut microbes were also recognized by anti-xylan and anti-xyloglucan antibodies (25% binding and 32% binding, respectively) compared to isotype controls **(Fig. 5e)**. Together, these results indicate that microbes with fiber glycans on their surfaces are present in the communities that sensitize the host to produce anti-fiber antibodies.

## Discussion

IgA secreted into the intestine functions in part by binding luminal microbes and preventing their adhesion to or translocation across the intestinal epithelium, thereby maintaining mucosal compartmentalization^30–32^. During homeostasis, anti-fiber IgA could similarly restrict the translocation of dietary glycans and glycan-coated microbes, but these antibodies also have the potential to shape the microbial ecosystem by blocking access to certain nutrients. While a secreted antibody targeting a bacterial protein has the potential to interfere with a limited set of related species (such as in the case of the fructan polysaccharide utilization locus of *Bacteroides thetaiotaomicron* VPI-5482 ^33^) a single anti-fiber IgA specificity could conceivably impact fiber harvest by many unrelated microbial taxa. The detection of IgA on gut bacteria (often referred to as “mFLOW” or “BugFACS”) is an increasingly standard assessment of the intestinal immune response. Our data indicate that IgA-positive bacteria identified using these tools include not only those that express targeted bacterial antigens but also those coated in dietary glycan antigens. Future studies will benefit from incorporating the contribution of anti-fiber IgA into the interpretation of IgA binding data. The ability of certain bacterial species to induce IgA responses to exogenous compounds has implications for the development of mucosal vaccines.

Fiber-specific IgM may function similarly to circulating antibodies that bind gut microbial surface structures and limit their dissemination during infection^34,35^. In cases of barrier disruption, anti-fiber antibodies could enable capture and degradation of fiber-coated microbes by immune cells to prevent their accumulation in the lamina propria. In other contexts, anti-fiber IgM could cause complement activation leading to inflammation. Previous work found that antibodies against *Streptococcus pneumonia* serotype 10A cross-react with plant pectin by virtue of a shared glycan epitope (beta-1,6-galactose)^36^. Although many of the polysaccharides recognized by the antibodies we report are unique to plants, this raises the prospect that gut bacterial glycans could mimic these structures. These results highlight the need to further understand the interactions between dietary fiber, the microbiota, and the immune system, especially when considering the immense diversity of dietary glycans in human diets. Our approach using libraries of polysaccharide-coated beads will enable future studies investigating humoral responses against pathogen and vaccine antigens to assess the potential for related epitopes present in the diet.

Microbes synthesize surface proteins and carbohydrates that elicit antibody production. For example, in *Bacteroides fragilis*, genetic deletion of a locus necessary for colonization diminishes IgA coating^37^, while in *Bacteroides thetaiotaomicron*, distinct capsular polysaccharides modulate antibody recognition and colonization^38^. In addition, *Akkermansia*, *E.coli*, and *Lachnospiraceae* strains also induce microbe specific responses in mice^39–41^. These findings implicate separate microbial ligands for host pattern recognition receptors in the induction of antibodies against these structures. Our results demonstrating physical associations between dietary glycans and microbial surfaces, as well as rescue of anti-fiber antibody production in germ-free mice colonized with microbes coated in fiber glycans, support the notion that bacterial binding to fiber enhances the host response.

Microbes in the gut lumen are monitored through well-described antigen-sampling pathways, including M-cell transcytosis, dendritic cell uptake, and secretory cell antigen passages^42^. Although there are no established analogous translocation pathways for fiber, it is possible that fibers present on microbial surfaces enter through one or more of these routes. Previous studies have detected unelicited natural antibodies that recognize herbal polysaccharides^43^ and total IgA can be increased by fiber supplimentation^44,45^. However, it is unclear how high molecular weight plant polysaccharides gain access to glycan-specific B cells. Several factors could influence whether B cells are exposed to fiber glycans, including structural features, lumenal environment (eg. mucus thickness)^46^, and microbial degradation^39–41,47^. We observed that both poorly fermentable (psyllium) and readily fermentable (karaya and galactomannan) fiber-types induce antibodies, suggesting that fermentability alone does not dictate antibody induction. However, the precise contributions of these factors remain unknown, as does their applicability to different fiber classes

Dietary fiber is widely recognized as a key contributor to human health and has been shown to modulate immune responses and influence outcomes across several disease contexts^48–50^. Therapeutic diets with varying amounts of fiber have shown promise as a treatment for patients with inflammatory bowel disease^50,51^. However, the immense variety of glycan structures present across distinct tissues in each plant, and across plant species, has hampered progress in determining which fiber types are beneficial and in which individuals^52,53^. It is posited that these therapeutic diets act by altering metabolism in fiber-consuming microbes, but this mechanism has not been definitively confirmed. Furthermore, some dietary interventions appear to exert opposing effects on disease symptoms versus markers of inflammation^51^. Our findings suggest that anti-psyllium antibodies are present in previous psyllium feeding studies^18,23^, although their contribution to the observed protection remains to be determined. Fulfilling the promise of personalized therapeutic fiber administration requires a deeper understanding of how plant glycans in fiber alter the immune system as well as the gut microbiota, both in healthy populations and in cases where the epithelial barrier is compromised^54,55^.

## Acknowledgements

We thank Jennifer K. Bando and members of the Patnode Lab for reviewing the manuscript and providing valuable feedback.

## Author Contributions

G.V. and M.L.P. planned and designed the experiments. M.L.P., A.H., and A.R.W. supervised experiments and obtained funding. G.V., C.A., M.E.G., N.H.K., M.L.P, K.R.I., and A.R. conducted the experiments. E.A.D., J.A., R.H., and A.H. provided human serum. G.V. and M.L.P. performed data analysis. G.V. and M.L.P. wrote the manuscript with assistance from C.A., A.R.W., E.A.D., A.H. All authors read and approved the manuscript.

## Competing Interests

The authors declare no competing interests.

**Correspondence and requests for materials** should be addressed to Michael L. Patnode.

## METHODS

### Mice

All experiments involving mice were carried out in accordance with protocols approved by the Animal Studies Committee of the University of California, Santa Cruz and the University of California, Berkeley. C57BL/6 and Swiss Webster mice (6-16 weeks) were reared and maintained in specific pathogen-free (SPF) facilities or, for gnotobiotic mice, maintained in cages located within flexible plastic isolators in the UCSC gnotobiotic facility or at UCB on an Innovive Innorack IVC Mouse 3.5 using Innovive’s caging (M-BTM bottom, MVX3 containment lid). Cages contained paper towels for environmental enrichment. For gnotobiotic colonization, pre-colonization fecal pellets were collected and cultured to verify the sterility of the mice. Female and male mice were used, and no sex differences were observed in the reported measures. T cell-deficient mice (B6.129P2-*Tcrb^tm1Mom^ Tcrd^tm1Mom^*/J) were obtained from Jackson Laboratories (002122). All mice were maintained on a strict light cycle (lights on at 0600h, off at 1800h).

### Human Samples

Deidentified human serum samples were obtained from healthy adult and child participants enrolled in a collaborative study conducted at the International Centre for Diarrhoeal Disease Research, Bangladesh (icddr,b). The cohort comprised adults (21-41 years old) and children (3-4 years old) with no major gastrointestinal surgery, recurrent intestinal inflammation, recent COVID-19 infection, recent antibiotic use, recent diarrhea, or other recent vaccination. Serum samples were collected at enrollment and again at 1-week post-enrollment. Following collection, serum was processed according to standard clinical procedures, stored at -80°C, and maintained as deidentified specimens until downstream analysis.

### Mouse feeding experiments

Dietary fibers (psyllium, karaya gum, and galactomannan) were either included at 9.2% w/w in a control diet during pelleting (Envigo, TD.170694 and Envigo, TD.250449) or mixed at 9.2% w/w with a powdered control diet (Envigo, T.2020XM) in collection cups (Avantor Science Central, 89508-716) or autoclavable bags (Fisher Scientific, 01-812-54). The prepared diets were then autoclaved for sterilization. On day 0, mice were switched from a pellet diet to the experimental diets for 11 or 12 days depending on experimental design. Powdered diets were hydrated with 100 mL of sterile water, and the resulting paste was pressed into a feeding dish and placed on the cage floor as previously described^11^. Food levels were monitored daily, and a freshly hydrated serving of the diet was supplied every 2 days. For experiments using antibiotics, ampicillin (1 g/L) and neomycin (0.5 g/L) were administered through the drinking water of mice 4 days before the initiation of experimental diets and continued for the duration of the experiment. Mice were switched to a low-fiber diet (Envigo, TD.130343) where indicated for 5 days.

### Bacterial samples for mouse colonization

For preparing the complete mouse microbiota culture collection, cecal contents and fecal pellets were harvested from two separate mice, each eating a low-fiber diet for 15 weeks to limit potential strain candidates. Samples were added to 5 mL of filtered TYG supp (supplemented tryptone yeast glucose) media^56^, mashed, filtered through a 100 µm cell strainer in an anaerobic chamber, and rinsed with 5 mL of filtered TYG. Filtrate (1 mL) was plated onto aerobic and anaerobic brain-heart-infusion (BHI) blood agar plates and allowed to grow at 37°C. After 5 days, the colonies were scraped and mixed in 5 mL of filtered TYG. The samples were then prepared in 1 mL aliquots for colonization. For colonization with clonal strains, aliquots of bacterial stocks were thawed, and the outer surfaces were sterilized with activated Clidox (Fisher Scientific, NC9122380 and NC0089321) before transferring into gnotobiotic isolators. Cell suspension (200 uL) was administered through a plastic-tipped oral gavage needle. Equal volumes of bacterial stocks were combined when colonizing germ-free mice with multiple bacterial strains. The colonized mice were bred and maintained in plastic gnotobiotic isolators. Fecal pellets from the progeny were collected and filtered (40 µm) using 5 mL of sterile water. Filtrate (200 uL) was administered to age-matched gnotobiotic mice through a plastic-tipped oral gavage needle. Identities of commensal gut bacterial isolates, *Bacteroides ovatus* ATCC-8483 (ATCC), *Bacteroides ovatus* WH514^57^, *Akkermansia muciniphila* NSD001 (Jax C57BL/6 microbiota), and *Escherichia coli* TSDC17.2^58,59^ were verified by full-length 16S rDNA sequencing. Each strain of the OMM12 community^60^ was grown according to the specifications provided by DSMZ, and pooled at equal OD600 for oral gavage of mice. C57BL/6NTac germ-free mice (Taconic Biosciences) were colonized with 200 µL of the OMM12 cocktail once via oral gavage as above. One week post-gavage, fecal samples were collected to confirm colonization using 16S amplicon sequencing.

### Mouse sample collections

Blood was collected from live mice via submandibular vein puncture into serum separator tubes (Fisher Scientific, 02-675-185). Blood samples were allowed to clot at room temperature for 30 mins and then centrifuged at 18213xg speed for 5 mins at 4°C to isolate the serum. After centrifugation, sodium azide (0.05% v/v) was added and the serum stored at -80°C until analysis. Mice were euthanized individually by CO□ asphyxiation followed by bilateral thoracotomy. Following euthanasia, whole blood was collected by cardiac puncture using a 22-gauge needle and processed in the same manner as described above. The cecum was collected, and a small incision was made in the cecal wall. Cecal contents were squeezed from the incision onto a sterile surface. The circular end of a sterile yellow plastic inoculation loop (Fisher Scientific, 22363600) was used to collect cecal contents (∼3 loops per mouse). Each loop was immediately flash-frozen in liquid nitrogen and transferred into a screw-cap tube. Samples were stored at -80°C until processing. For cecal content processing, a freshly made working solution (10 mL) of phosphate-buffered saline containing 0.01% sodium azide and one protease inhibitor tablet (Sigma, 11836170001) was prepared (PBS-SA-PI). Frozen cecal contents were weighed, and PBS-SA-PI was added (3 µL/mg of cecal contents). A sterile 3/8-inch stainless steel grinding ball (McMaster-Carr, 96455k73) was added to each tube. Samples were homogenized using a bead beater (Biospec, 1001) at 14000 rpm for 1 min, then centrifuged at 18213xg speed for 5 mins at 4°C. The resulting supernatant was transferred to a screw-cap tube, and additional sodium azide was added to a final concentration of 0.05% (v/v). Processed samples were stored at -80°C until further analysis.

### Fiber glycan bead library

Biotinylated glycans were generated as described previously^11^. Briefly, glycan preparations were suspended in water at a concentration of 20 mg/mL, heated by sonicating for 1 min, and then centrifuged at 18213xg speed for 10 min. The resulting solutions were diluted 1:4 in water (v/v). TFPA-PEG3-biotin (Thermofisher, 21303), dissolved in DMSO at 10 mg/mL, was added to the glycan solutions at a ratio of 1:5 (v/v). Samples were subjected to UV irradiation for 10 mins (Fisher Scientific, 13-245-221) and then diluted 1:4 (v/v) to facilitate desalting using 96-well ZEBA plates with a 7kDa molecular weight cut-off (Thermo Scientific, 89808).

Biotinylated glycans were incubated with paramagnetic, streptavidin-coated silica beads (Millipore Sigma, LSKMAGT) at a 1:2 v/v ratio for 1 hr at room temperature. Beads were washed three times with 150 µL HNTB buffer (10 mM HEPES, 150 mM NaCl, 0.05% Tween-20, 0.1% bovine serum albumin) using a magnetic stand. Beads were subsequently incubated for 30 mins with 1 µg/mL streptavidin-fluorophore mixtures in HNTB. The process of washing, biotin-glycan incubation, and re-washing was repeated for one additional cycle. No impact of streptavidin-fluorophore selection on antibody binding was detected after permutation of color combinations in replicate bead libraries.

### Anti-fiber antibody detection assay

An aliquot of the pooled bead library was added to the wells of a 96-well plate. A 10-15 µL sample of monoclonal antibody (CarboSource, SKCMA-AM1 and SKCMA-AM2), serum, cecal contents, or colostrum was added to the beads and incubated at 4°C with constant rotation for 1 hr. The beads were then collected using a magnetic rack and washed three times with 150 µL HNTB. Beads were incubated with the appropriate fluorescent secondary antibodies (1 µg/mL) at 4°C with constant rotation for 30 mins and washed three times with 150 µL HNTB. Beads were run on an LSR II (BD Biosciences), CytoFLEX (Beckman Coulter), or Attune (Thermo Fisher Scientific). The beads were identified based on streptavidin fluorescence, and fluorescence of the secondary antibody was measured for each bead type. Separate aliquots of the bead library were incubated with fluorescent secondary antibodies in the absence of primary antibodies to establish background fluorescence levels.

### ELISA

Carbohydrates were diluted with sterile water (1:1000) and 60 µL was plated onto Immulon 2HB 96-well plates (Thermo Fisher Scientific, 3455). The plates were left at 37°C overnight, uncovered to induce complete evaporation. The following day, the plates were washed with 200 µL PBS containing 1% Tween-20 (PBS-T) and blocked for 1 hr at 4°C with 200 µL/well blocking buffer (PBS containing 1-5% bovine serum albumin). The plates were then washed three times with 200 µL PBS-T and incubated with 50 µL diluted serum (1:50, 1:100, or 1:200 with PBS-T) or cecal contents (1:10 or 1:25 with PBS-T) for 1 hr at 4°C. For detection, plates were washed three times with 200 µL PBS-T, and bound antibodies were detected using 50 µL of the appropriate horseradish peroxidase (HRP)-conjugated secondary antibodies (1 µg/mL) and incubated for 30 mins at 4°C. After washing three times with 200 µL PBS-T, 50 µL/well of 3,3′,5,5′-tetramethylbenzidine (TMB) substrate (Thermo Fisher Scientific, N301) was added, and the reaction was developed for 5-15 mins in the dark. The reaction was stopped with 50 µL/well of 0.2 M sulfuric acid, and the absorbance was read at 450 nm and 570 nm using a microplate reader.

Unconjugated capture antibodies were diluted to 0.5 µg/mL with PBS and plated onto Immulon 2HB 96-well plate(s) (Thermo Fisher Scientific, 3455) and incubated overnight at 4°C with 60 µL/well. The following day, the plates were blocked as described above. The plates were then washed three times with 200 µL PBS-T, and samples were diluted with PBS-T and added at 50 µL/well. A standard curve was prepared using serial dilutions (1:2) of purified antibodies of the same isotype as the target, starting at 2 µg/mL or 0.5 µg/mL. Plates were incubated for 1 hr at 4°C, followed by three washes with 200 µL PBS-T. The plates were then processed for detection as described above.

### Microbial flow cytometry

Fresh fecal pellets were collected from mice and weighed. Fecal pellets received 1 mL of PBS containing 1% bovine serum albumin and 0.05% sodium azide (BS-PBS). A sterile 3/8-inch stainless steel grinding ball (McMaster-Carr, 96455k73) was placed into each tube, and samples were homogenized using a bead beater (Biospec, 1001) at 1,400□ rpm for 1 min. Following homogenization, the homogenized samples were filtered twice through a 70 µm nylon mesh filter to remove debris. Filtrates were then transferred to a U-bottom 96-well plate with volumes adjusted based on sample weights to achieve a final concentration of 0.5-1 mg of fecal material per well. Bacterial pellets were washed twice by centrifugation at 3,220xg for 5□ mins at 4□ °C. Where indicated, bacterial pellets were incubated with 50 µL BS-PBS, diluted xylanase (1 U/mL with BS-PBS [Neogen, E-XYLNP]), or diluted xyloglucanase (1 U/mL with BS-PBS [Neogen, E-XEGP]) for 15 mins at 37□ °C on a plate shaker at 250 rpm (SCILOGEX, 822000049999) and then washed twice by centrifugation at 3,220xg for 5 mins at 4°C. Bacterial pellets were resuspended with 20 µL of diluted monoclonal antibodies (1 µg/mL with BS-PBS) or the respective isotype control (1 µg/mL) and incubated for 1 hr at 4°C on a plate shaker at 250 rpm. Following incubation, samples were washed twice with 200 µL BS-PBS by centrifugation (3,220xg, 5□ mins, 4□ °C). 50 µL of fluorescent secondary antibodies (1 µg/mL) stained bacterial pellets for 30 mins at 4 °C with shaking. After washing twice, the samples were stained with 50µL of 10 µM Syto62 (Thermo Fisher Scientific, S11344) and incubated for 5 mins at room temperature in the dark with shaking. Samples were washed twice and resuspended with a final volume of 150 µL BS-PBS. The final suspensions were passed through a 70 µm filter into a U-bottom plate to remove aggregates.

### V4 16S rDNA sequencing

Fecal DNA was isolated using the DNeasy PowerSoil Pro Kit (Qiagen, 47014). Extracted DNA was quantified using Qubit, normalized, and the V4 region was PCR amplified. Each V4 amplicon was annealed to a sample-specific i5 and i7 Illumina adapter index. The samples were pooled at equal concentration and run on a 2×250 MiSeq Illumina instrument. Reads were demultiplexed and analyzed using QIIME2 (version 2021.4). Sequences were singlet filtered and used to generate amplicon sequence variants (ASVs). The taxonomy was assigned by the Silva 138 classifier for the 515F/806R region (V4). These results were then summarized to produce sequence counts and the relative abundance of taxa per sample.

## Statistics and Reproducibility

Statistical analysis was performed in GraphPad Prism 11 and R (v4.5.2) using RStudio (v2026.01.0+392). All glycan bead flow cytometry data is presented in supplementary tables including empty bead values, which were averaged and used for background subtraction unless otherwise indicated. For statistical testing, bead types were filtered for those where the raw signal was significantly greater than that of empty beads (Paired t-test and Benjamini-Hochberg (BH) FDR p value correction, q < 0.1) and exceeded a minimum fold change threshold (FC >= 1.5) in at least one of the two groups being compared. After filtering, differences between groups were calculated using a Student’s t test and BH FDR p value correction (q < 0.05) and exceeding a minimum fold change threshold (FC >= 1.5 for mouse samples and FC >= 1.25 for human samples), unless otherwise indicated. For analysis of correlations between antibody responses to glycans in human samples, a Spearman’s rank correlation coefficient was calculated for each glycan-glycan pair across individuals. For anti-fiber bar plots, antibody binding to bacteria, and ELISAs, significance was determined using Student’s t test. For anti-psyllium antibody comparisons collected from mice at different time points, significance was determined using a paired t-test. All t-tests were two-tailed with a p-value cutoff of 0.05. Mean values and standard deviations or 95% confidence intervals are shown in the figures as indicated.

## Data Availability

Raw sequencing data have been uploaded to ENA: Accession number PRJEB115493.

## Extended Data Figure and Table Legends

**Extended Data Fig. 1.**
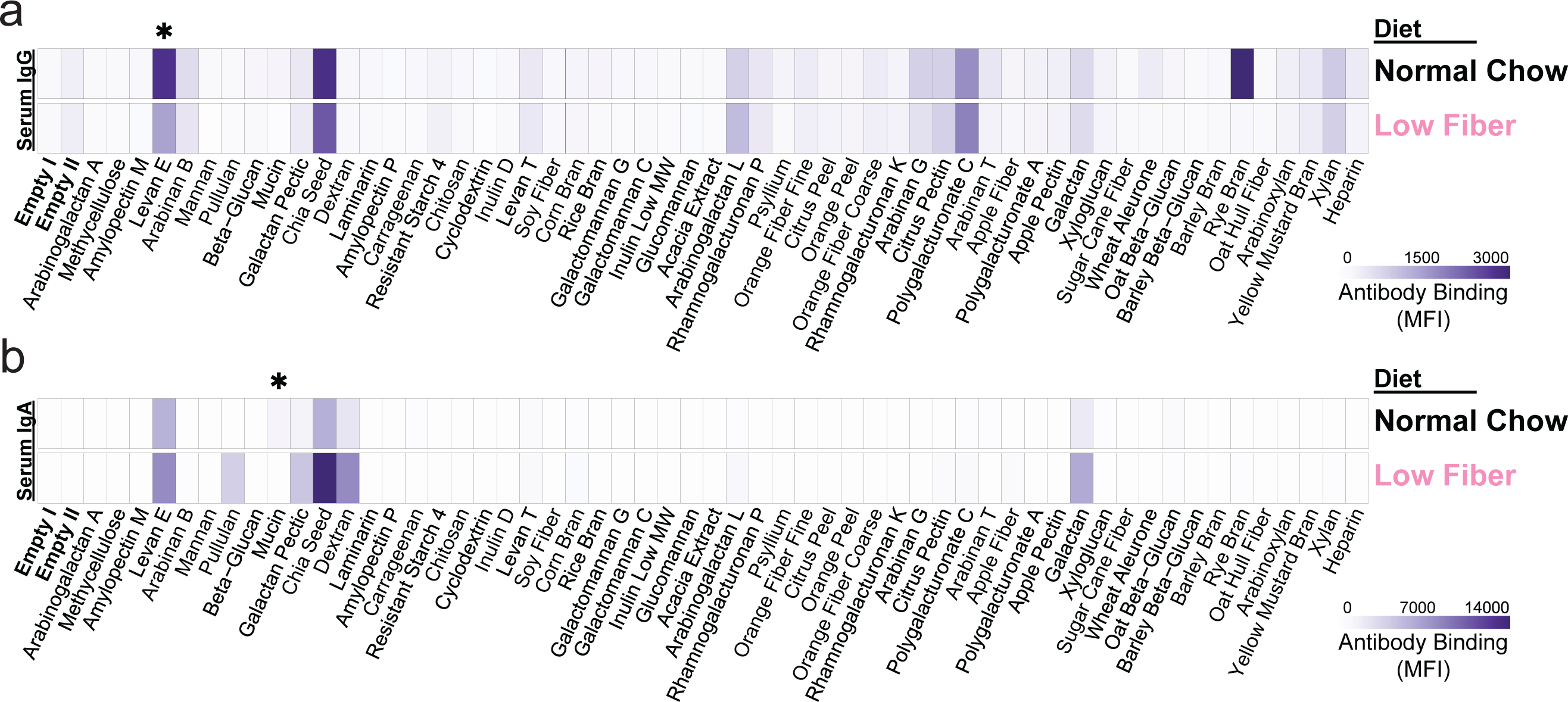
Mice produce circulating anti-fiber IgG and IgA. (a and b) Anti-fiber IgG (a) and IgA (b) was measured in serum from C57BL/6 mice eating normal chow or a low fiber diet by flow cytometry using a glycan-coated bead library. Each row shows the average of 7-8 mice. Results are representative of 2 independent experiments. Heatmap values are corrected by subtracting the average of 3 secondary-only controls and the average of 2 empty bead populations (*, Student’s t test, q < 0.05, FC >= 1.5).

**Extended Data Fig. 2.**
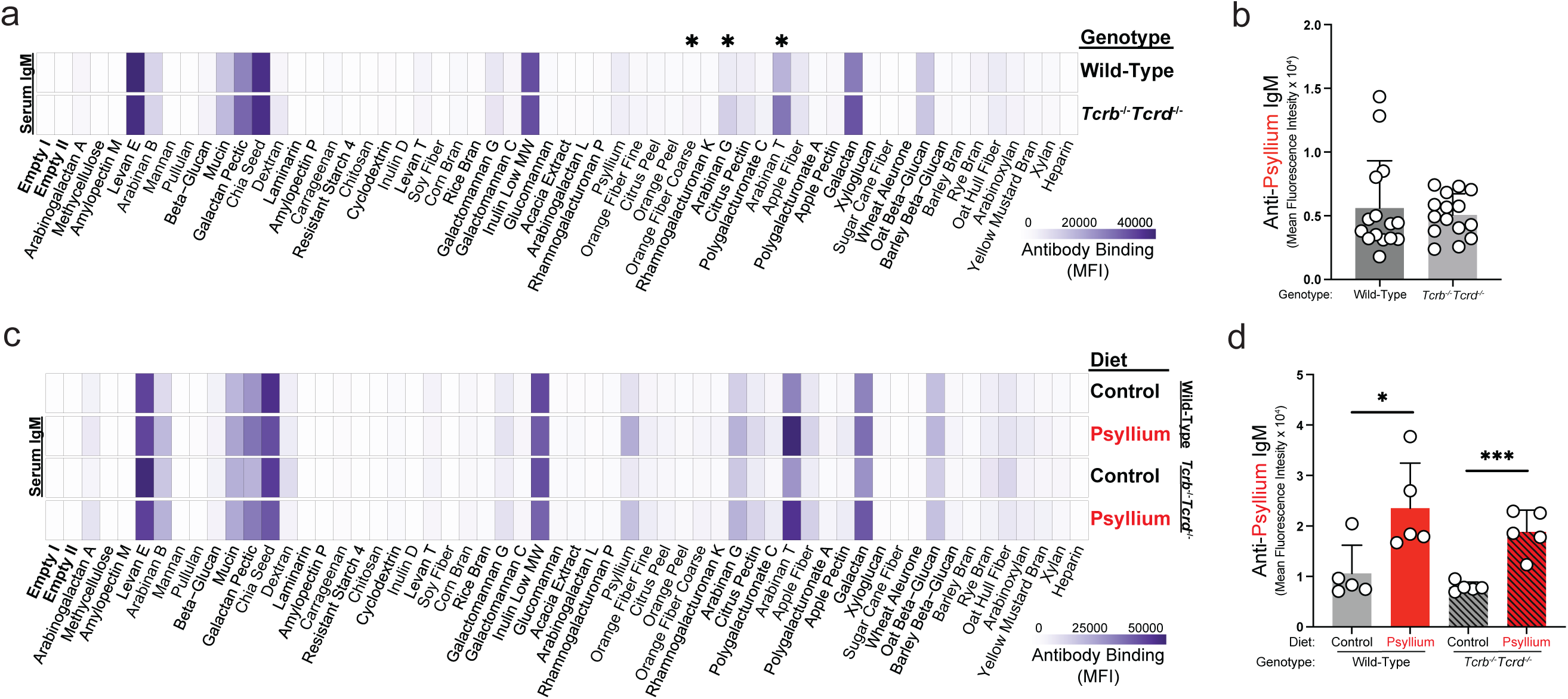
Induction of anti-psyllium antibodies does not require T cells. (a) Serum from wild-type or *Tcrb^-/-^Tcrd^-/-^* C57BL/6 mice collected fed normal chow (day -2) was tested using a glycan-coated bead library to measure baseline anti-fiber IgM differences (*, Student’s t test, q < 0.05, FC >= 1.5). Each row shows the average of 15 mice. (b) Individual values for anti-psyllium IgM in serum on day -2. (c) Anti-fiber IgM in serum on day 17 after eating a control or 9.2% w/w psyllium-supplemented diet for 12 days followed by a low fiber diet for 5 days. Each row shows the average of 5 mice. Results are representative of 3 independent experiments. (d) Individual values for anti-psyllium IgM in serum on day 17 (Student’s t test, *p < 0.05 and ***p < 0.001). Heatmap values are corrected by subtracting the average of 3 secondary-only controls and the average of 2 empty bead populations. Bars, mean + SD.

**Extended Data Fig. 3.**
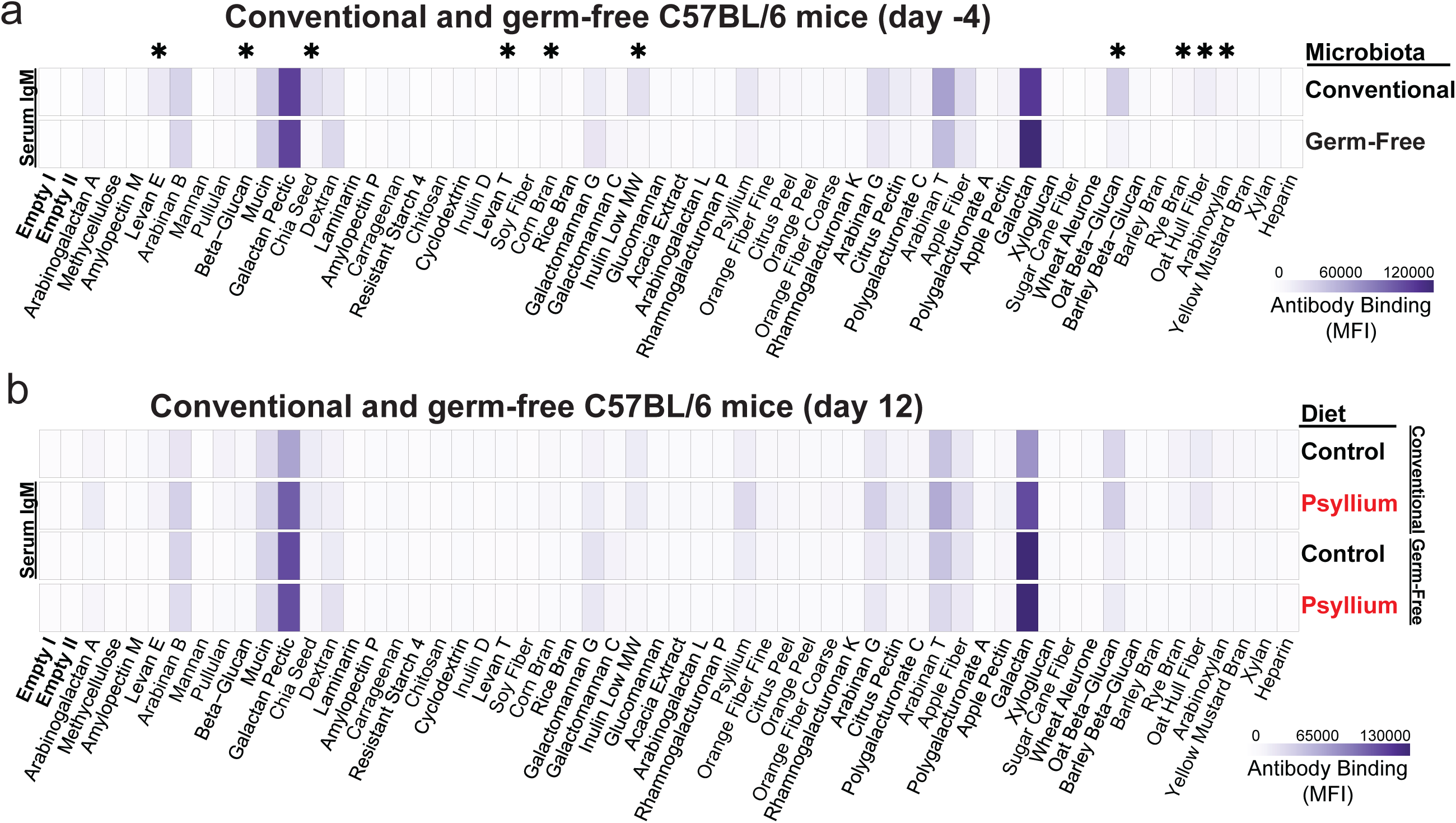
Absence of anti-fiber response in Germ-free mice. (a and b) Circulating anti-fiber IgM from conventional or germ-free C57BL/6 mice before (a, day -4) and after eating a control or 9.2% w/w psyllium-supplemented diet for 12 days (b) was tested using a glycan-coated bead library. Each row shows the average of 10-12 mice (a) or 5-6 mice (b) (*, Student’s t test, q < 0.05, FC >= 1.5). Results are representative of >3 independent experiments. Heatmap values are corrected by subtracting the average of 3 secondary-only controls and the average of 2 empty bead populations.

**Extended Data Fig. 4.**
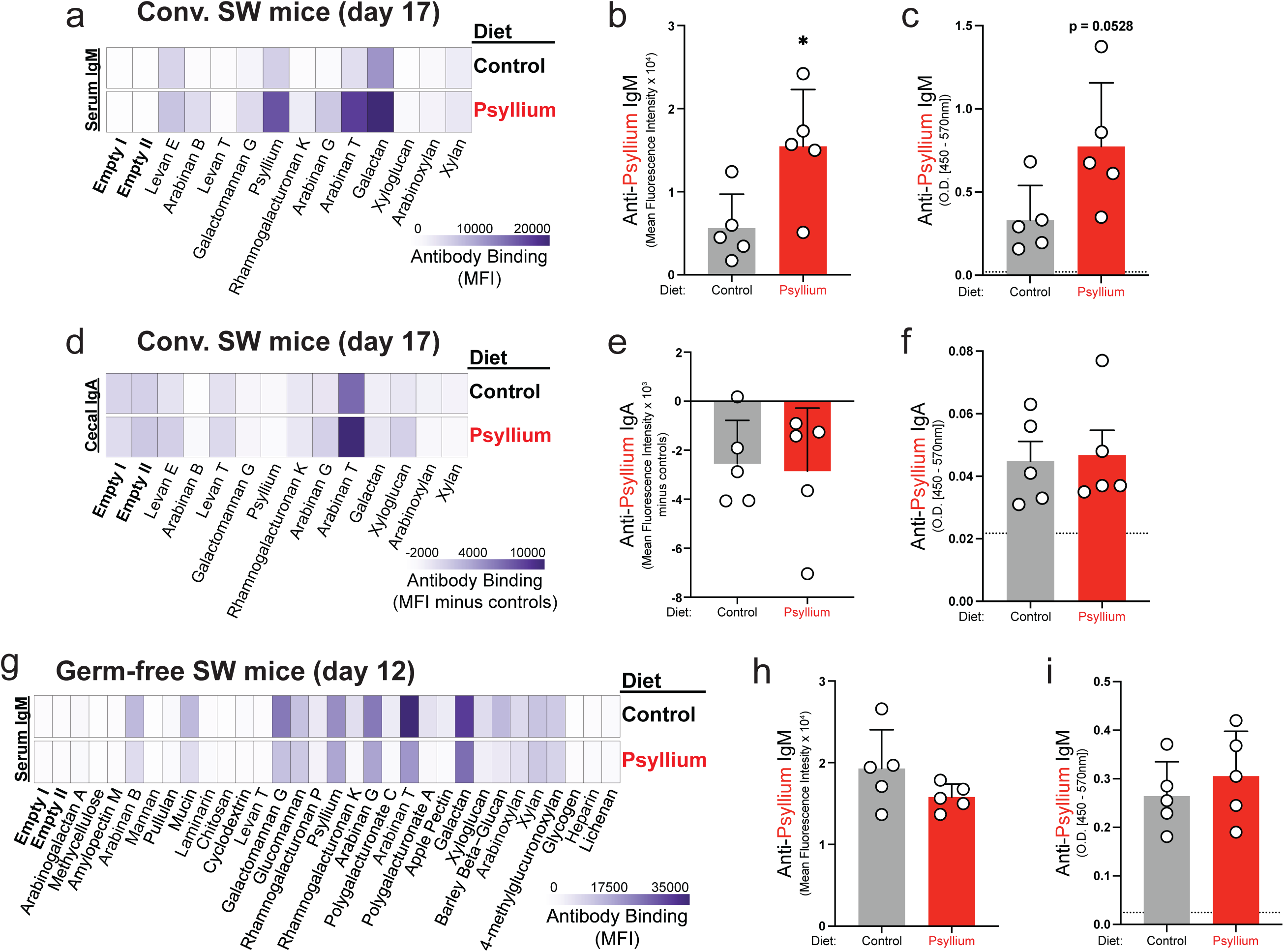
Circulating response against psyllium in conventionally-raised Swiss Webster mice. (a) Circulating anti-fiber IgM from conventional Swiss Webster (SW) mice eating a control or 9.2% w/w psyllium-supplemented diet for 12 days followed by a low fiber diet for 5 days was tested using a glycan-coated bead library. Each row shows the average of 5 mice. (b) Individual values for anti-psyllium IgM in serum on day 17 (Student’s t test, *p < 0.05). (c) ELISA for anti-psyllium IgM in serum on day 17. (d) Bead library showing cecal anti-fiber IgA signals. Each row shows the average of 5 mice. (e) Individual values for anti-psyllium IgA in cecal contents on day 17. (f) ELISA for anti-psyllium IgA in cecal contents on day 17. (g) Circulating anti-fiber IgM from germ-free SW mice eating a control or 9.2% w/w psyllium-supplemented diet for 12 days was tested using a glycan-coated bead library. Each row shows the average of 5 mice. (h) Individual values for anti-psyllium IgM in serum on day 12. (i) ELISA for anti-psyllium IgM in serum on day 12. Heatmap values are corrected by subtracting the average of 3 secondary-only controls and the average of 2 empty bead populations. Dotted lines, background signal (average of 3 secondary-only controls); bars, mean + SD.

**Extended Data Fig. 5.**
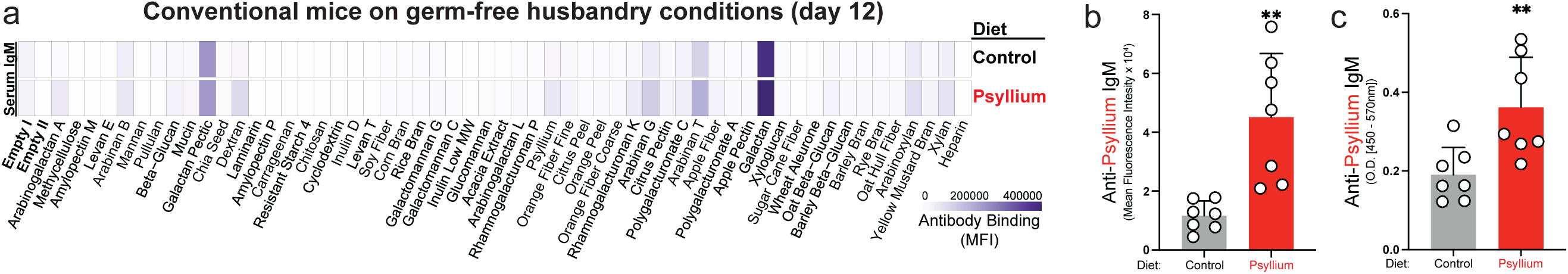
Germ-free husbandry conditions do not inhibit the induction of anti-psyllium antibodies. (a) Circulating anti-fiber IgM from conventionally raised C57BL/6 mice (raised from birth using the same autoclave-sterilized bedding and chow used for germ-free mice) eating a control or 9.2% w/w psyllium-supplemented diet for 12 days was tested using a glycan-coated bead library. Each row represents the average of 7 mice. (b) Individual values for anti-psyllium IgM in serum on day 12 (Student’s t test, **p < 0.01). (c) ELISA for anti-psyllium IgM in serum on day 12 (Student’s t test, **p < 0.01). Heatmap values are corrected by subtracting the average of 3 secondary-only controls and the average of 2 empty bead populations. Dotted lines, background signal (average of 3 secondary-only controls); bars, mean + SD.

**Extended Data Fig. 6.**
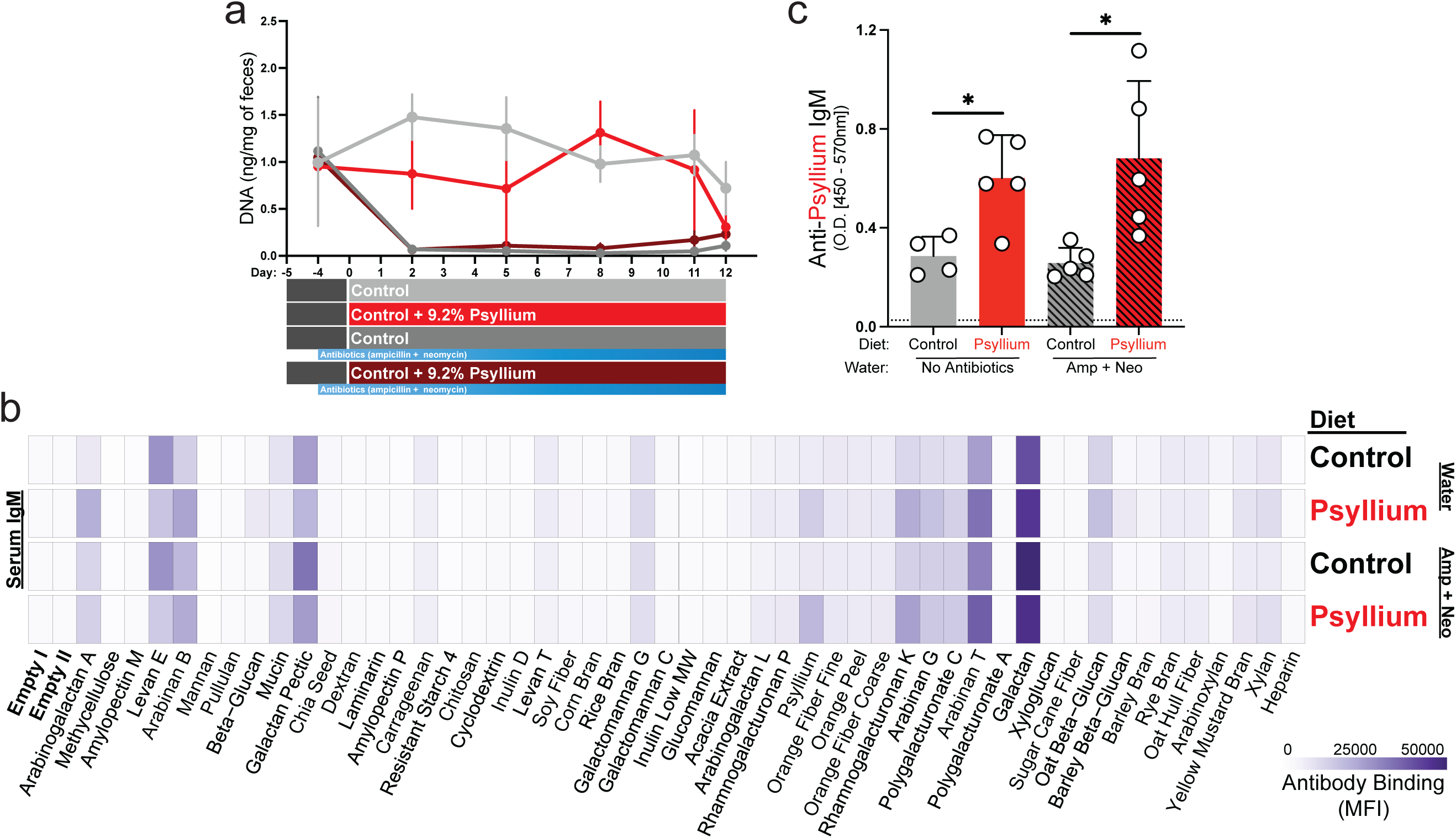
Antibiotic treatment does not inhibit the induction of circulating anti-psyllium IgM. (a) Fecal DNA from C57BL/6 mice with or without antibiotic treatment (eating a control or 9.2% w/w psyllium-supplemented diet for 12 days) was quantified before starting the fiber feedings (day -4) and throughout the feedings as a proxy for microbial abundance in the gut. (b) Circulating anti-fiber IgM was tested using a glycan-coated bead library. Each row represents the average of 5 mice. (c) ELISA for anti-psyllium IgM in serum on day 12 (Student’s t test, *p < 0.05). Heatmap values are corrected by subtracting the average of 3 secondary-only controls and the average of 2 empty bead populations. Dotted lines, background signal (average of 3 secondary-only controls); bars, mean + SD.

**Extended Data Fig. 7.**
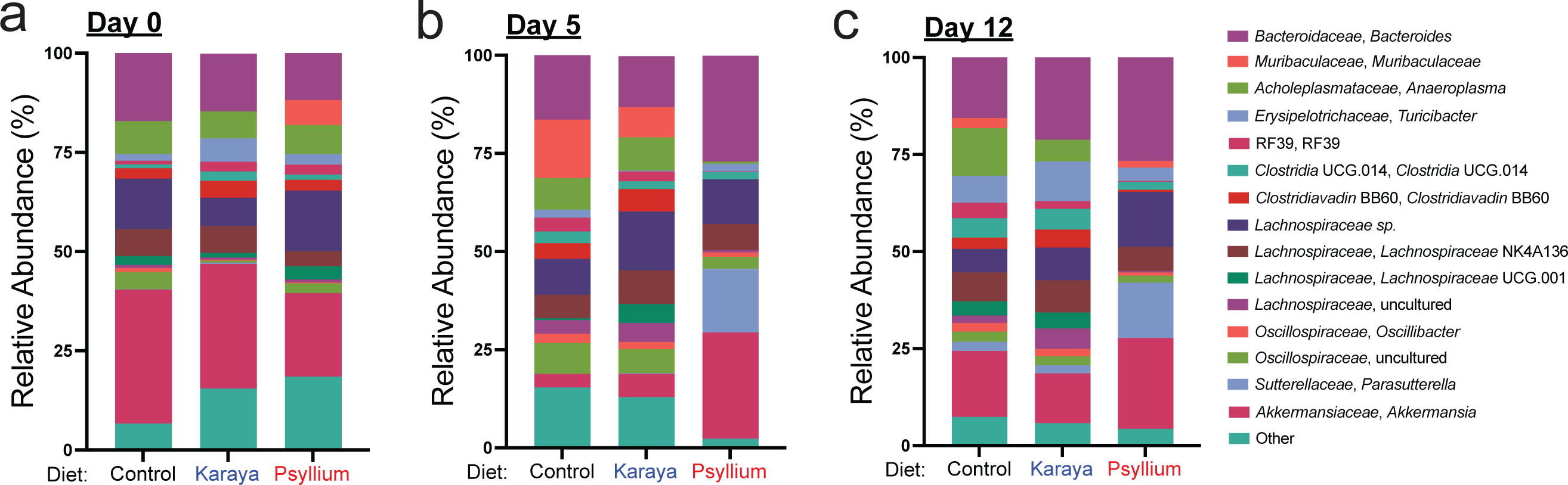
Psyllium feeding alters gut microbiota composition. (a - c) Relative abundance of bacterial taxa at the genus level in control, 9.2% w/w karaya, and 9.2% w/w psyllium fed mice at day 0 (a), day 5 (b), and day 12 (c) (control, n = 6; karaya, n = 6; psyllium, n = 6). The stacked bar plots display the top 15 most abundant taxa determined from day 12 sequences, with the remaining taxa classified as “other”.

**Extended Data Fig. 8.**
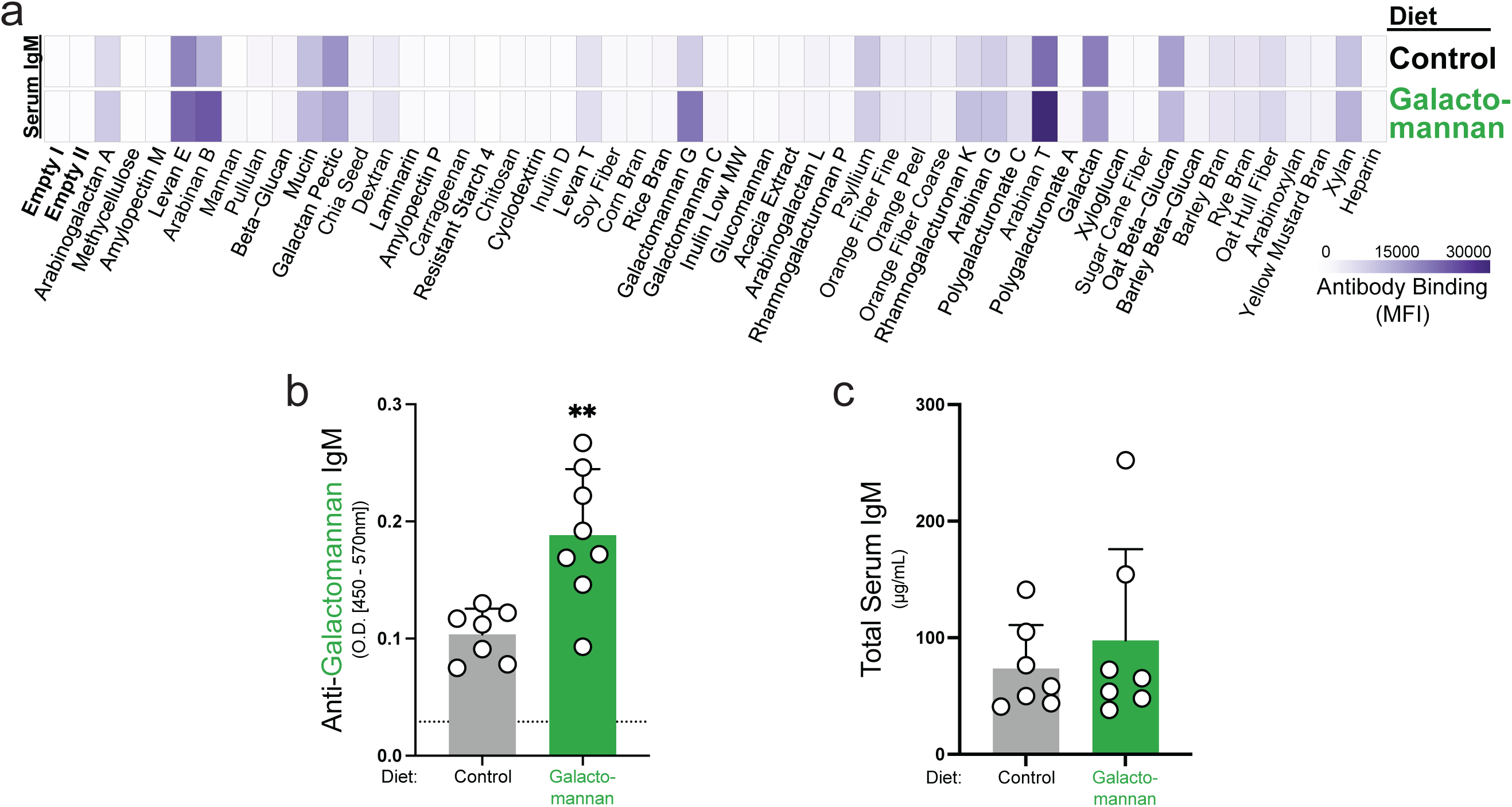
Galactomannan feeding induces circulating anti-galactomannan IgM. (a) Circulating anti-fiber IgM from C57BL/6 mice eating a control or 9.2% w/w galactomannan-supplemented diet for 11 days was tested using a glycan-coated bead library. Each row represents the average of 7-8 mice. Results are representative of >3 independent experiments. (b) ELISA for anti-galactomannan IgM in serum on day 11 (Student’s t test, **p < 0.01). (c) Quantification of total serum IgM on day 11. Heatmap values are corrected by subtracting the average of 3 secondary-only controls and the average of 2 empty bead populations. Dotted lines, background signal (average of 3 secondary-only controls); bars, mean + SD.

**Extended Data Fig. 9.**
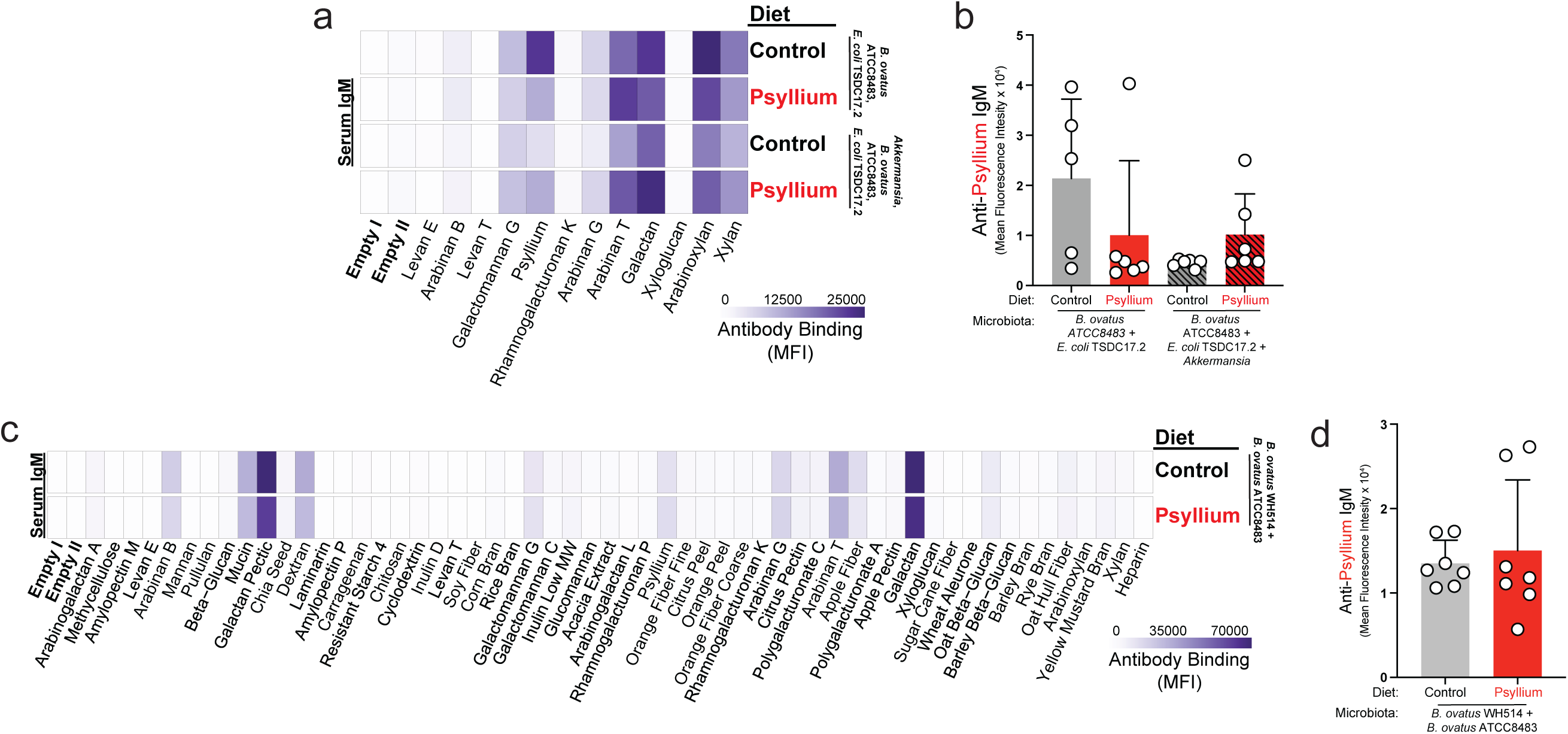
Diverse bacterial isolates are not sufficient for the induction of circulating anti-psyllium IgM. (a) Circulating anti-fiber IgM from germ-free Swiss Webster mice colonized with a 2 (*Bacteroides ovatus* ATCC8483 and *E. coli* TSDC17.2) or 3 (*B. ovatus* ATCC8483, *E. coli* TSDC17.2, and *Akkermansia muciniphila* NSD001) member community eating a control or 9.2% w/w psyllium-supplemented diet for 12 days was tested using a glycan-coated bead library. Each row represents the average of 5-6 mice. (b) Individual values for anti-psyllium IgM in serum on day 12. (c) Circulating anti-fiber IgM from germ-free C57BL/6 mice colonized with a 2-member community (*B. ovatus* WH514 and *B. ovatus* ATCC8483) eating a control or 9.2% w/w psyllium-supplemented diet for 12 days was tested using a glycan-coated bead library. Each row represents the average of 7 mice. (d) Individual values for anti-psyllium IgM in serum on day 12. Heatmap values are corrected by subtracting the average of 3 secondary-only controls and the average of 2 empty bead populations. Bars, mean + SD.

**Extended Data Fig. 10.**
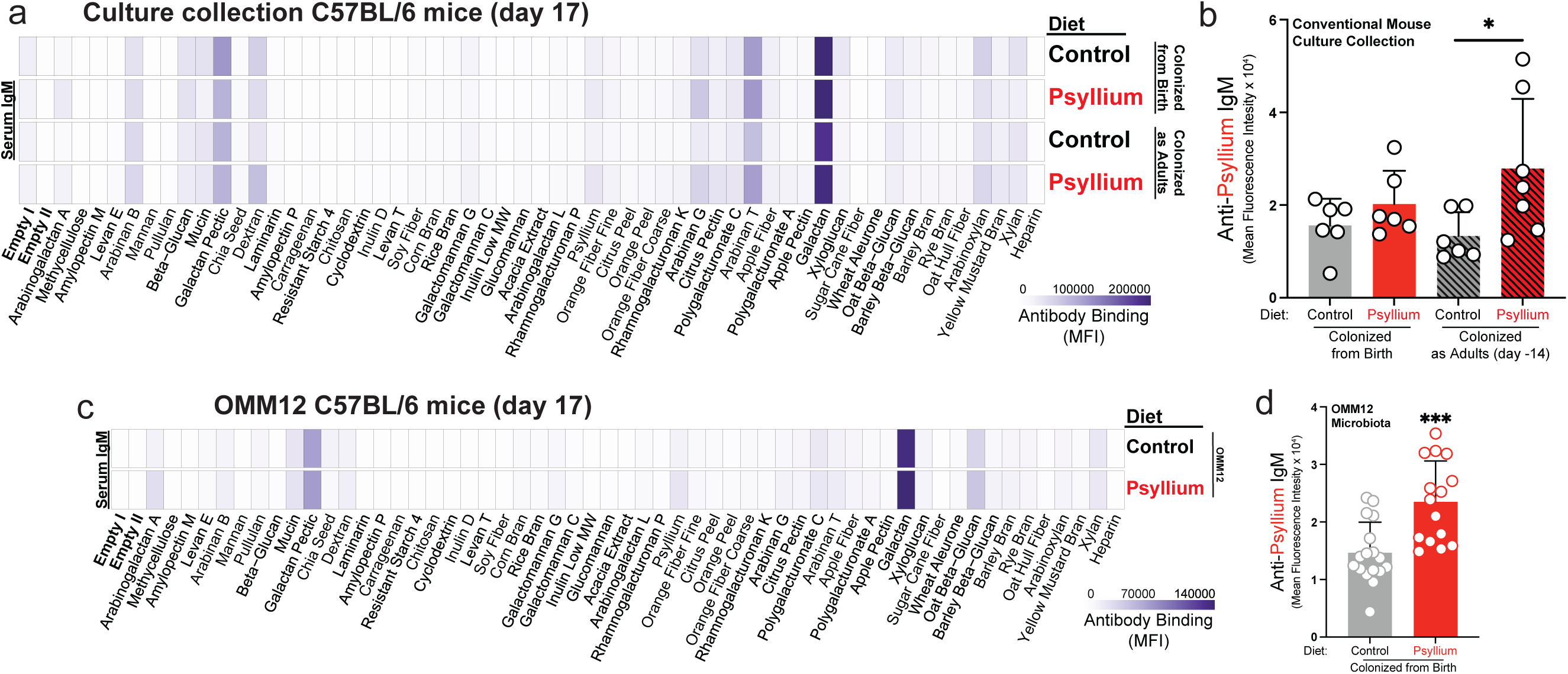
Culturable mouse microbiota enables germ-free mice to generate anti-psyllium antibodies. (a) Circulating anti-fiber IgM from germ-free C57BL/6 mice colonized with a conventionally raised mouse culture collection from birth or as adults eating a control or 9.2% w/w psyllium-supplemented diet for 12 days followed by a low fiber diet for 5 days was tested using a glycan-coated bead library. Each row represents the average of 6-7 mice. (b) Individual values for anti-psyllium IgM in serum on day 17 (Student’s t test, *p < 0.05). (c) Circulating anti-fiber IgM from germ-free C57BL/6 mice colonized with an OMM12 community eating a control or 9.2% w/w psyllium-supplemented diet for 12 days followed by a low fiber diet for 5 days was tested using a glycan-coated bead library. Each row represents the average of 15-17 mice. Results are representative of 2 independent experiments. (d) Individual values for anti-psyllium IgM in serum on day 17 (Student’s t test, ***p < 0.001). Heatmap values are corrected by subtracting the average of 3 secondary-only controls and the average of 2 empty bead populations. Bars, mean + SD or mean + 95% confidence interval (d).

**Supplemental Table 1. Anti-fiber antibodies in human samples.** (a) Geometric mean circulating IgM signal for each bead type at week 0 and week 1, and Spearman’s Correlation between timepoints. (b) Geometric mean of pooled colostrum IgA signal for each bead type.

**Supplemental Table 2. Anti-fiber antibodies in mice eating different diets.** (a) Empty-subtracted circulating anti-fiber IgM signal for each bead type in normal chow and low fiber-fed mice. (b) Empty-subtracted circulating anti-fiber IgM signal for each bead type in control and psyllium-fed mice. (c) Log2-transformed secreted anti-fiber IgA signal for each bead type in control and psyllium-fed mice.

**Supplemental Table 3. Circulating anti-fiber IgG and IgA in mice eating normal chow or a low fiber diet.** (a) Empty-subtracted circulating anti-fiber IgG signal for each bead type in normal chow and low fiber-fed mice. (b) Empty-subtracted circulating anti-fiber IgA signal for each bead type in normal chow and low fiber-fed mice.

**Supplemental Table 4. Circulating anti-fiber IgM in wild-type and *Tcrb*^-/-^*Tcrd*^-/-^ mice.** (a) Empty-subtracted circulating anti-fiber IgM signal for each bead type in wild-type and *Tcrb*^-/-^*Tcrd*^-/-^ mice at day -2. (b) Empty-subtracted circulating anti-fiber IgM signal for each bead type in wild-type and *Tcrb*^-/-^ *Tcrd*^-/-^ mice at day 17.

**Supplemental Table 5. Circulating anti-fiber IgM in conventional and germ-free C57BL/6 mice fed a control or psyllium diet.** (a) Empty-subtracted circulating anti-fiber IgM signal for each bead type in conventional and germ-free C57BL/6 mice at day -4. (b) Empty-subtracted circulating anti-fiber IgM signal for each bead type in conventional and germ-free C57BL/6 mice at day 12.

**Supplemental Table 6. Anti-fiber antibodies in conventional and germ-free Swiss Webster mice eating a control or psyllium diet.** (a) Empty-subtracted circulating anti-fiber IgM signal for each bead type in control and psyllium-fed conventional Swiss Webster mice. (b) Empty-subtracted secreted anti-fiber IgA signal for each bead type in control and psyllium-fed conventional Swiss Webster mice. (c) Empty-subtracted circulating anti-fiber IgM signal for each bead type in control and psyllium-fed germ-free Swiss Webster mice.

**Supplemental Table 7. Circulating anti-fiber IgM in mice fed a control or psyllium diet.** (a) Empty-subtracted circulating anti-fiber IgM signal for each bead type in conventional C57BL/6 mice maintained with germ-free husbandry conditions.

**Supplemental Table 8. Circulating anti-fiber IgM in antibiotic-treated and untreated mice.** (a) Empty-subtracted circulating anti-fiber IgM signal for each bead type in control and psyllium-fed mice with or without antibiotic treatment.

**Supplemental Table 9. Gut microbiota composition during fiber-feeding.** (a) Relative abundance (%) for each mouse across all included days. Taxa shown are the Top 15 ranked by mean abundance on Day 12 from all mice, with all remaining taxa collapsed into Other.

**Supplemental Table 10. Circulating anti-fiber IgM signals in mice fed a control or galactomannan diet.** (a) Empty-subtracted circulating anti-fiber IgM signal for each bead type in control and galactomannan-fed mice.

**Supplemental Table 11. Circulating anti-fiber IgM in mice colonized with minimal microbial communities fed a control or psyllium diet.** (a) Empty-subtracted circulating anti-fiber IgM signal for each bead type in control and psyllium-fed mice colonized with a 2-bacteria community (*Bacteroides ovatus* ATCC8483 + *E. coli* TSDC17.2) or 3-bacteria community (*B. ovatus* ATCC8483 + *E. coli* TSDC17.2 + *Akkermansia muciniphila*). (b) Empty-subtracted circulating anti-fiber IgM signal for each bead type in control and psyllium-fed mice colonized with a 2-bacteria community (*Bacteroides ovatus* ATCC8483 + WH514).

**Supplemental Table 12. Circulating anti-fiber IgM in mice colonized with communities of culturable isolates fed a control or psyllium diet.** (a) Empty-subtracted circulating anti-fiber IgM signal for each bead type in control and psyllium-fed mice colonized from birth or colonized as adults with a community of culturable isolates. (b) Empty-subtracted circulating anti-fiber IgM signal for each bead type in control and psyllium-fed mice colonized with an OMM12 model community from birth.

**Supplemental Table 13. Monoclonal antibody recognition of fiber-coated beads and fecal bacteria.** (a) Antibody isotype, antibody antigen, fecal bacteria binding values (percentage and MFI), and bead library binding values.

